# Increasing adult-born neurons protects mice from epilepsy

**DOI:** 10.1101/2023.07.08.548217

**Authors:** Swati Jain, John J. LaFrancois, Kasey Gerencer, Justin J. Botterill, Meghan Kennedy, Chiara Criscuolo, Helen E. Scharfman

## Abstract

Neurogenesis occurs in the adult brain in the hippocampal dentate gyrus, an area that contains neurons which are vulnerable to insults and injury, such as severe seizures. Previous studies showed that increasing adult neurogenesis reduced neuronal damage after these seizures. Because the damage typically is followed by chronic life-long seizures (epilepsy), we asked if increasing adult-born neurons would prevent epilepsy. Adult-born neurons were selectively increased by deleting the pro-apoptotic gene *Bax* from Nestin-expressing progenitors. Tamoxifen was administered at 6 weeks of age to conditionally delete *Bax* in Nestin-CreER^T2^*Bax*^fl/fl^ mice. Six weeks after tamoxifen administration, severe seizures (status epilepticus; SE) were induced by injection of the convulsant pilocarpine. After mice developed epilepsy, seizure frequency was quantified for 3 weeks. Mice with increased adult-born neurons exhibited fewer chronic seizures. Postictal depression was reduced also. These results were primarily in female mice, possibly because they were the more affected by *Bax* deletion than males, consistent with sex differences in *Bax*. The female mice with enhanced adult-born neurons also showed less neuronal loss of hilar mossy cells and hilar somatostatin-expressing neurons than wild type females or males, which is notable because these two hilar cell types are implicated in epileptogenesis. The results suggest that selective *Bax* deletion to increase adult-born neurons can reduce experimental epilepsy, and the effect shows a striking sex difference. The results are surprising in light of past studies showing that suppressing adult-born neurons can also reduce chronic seizures.

## INTRODUCTION

It has been shown that neurogenesis occurs in the hippocampal dentate gyrus (DG) during adult life of mammals (Taupin 2006; Gage et al. 2008; Altman 2011; Kempermann 2012; Kazanis 2013). It is important to note that this idea was challenged recently (Paredes et al. 2018; Sorrells et al. 2018) but afterwards more studies provided support for the original idea (Boldrini et al. 2018; Kempermann et al. 2018; Tartt et al. 2018; Moreno-Jimenez et al. 2019; Tobin et al. 2019).

In the DG, adult-born neurons are born in the subgranular zone (SGZ; Altman and Das 1965; Kaplan and Hinds 1977; Altman 2011). Upon maturation, newborn neurons migrate to the granule cell layer (GCL; Cameron et al. 1993), develop almost exclusively into GCs, and integrate into the DG circuitry like other GCs (Ramirez-Amaya et al. 2006; Kempermann et al. 2015).

Prior studies suggest that the immature adult-born GCs can inhibit the other GCs (Ash et al. 2023) especially when they are up to 6 weeks-old (Drew et al. 2016). By inhibition of the GC population, young adult-born GCs could support DG functions that require GCs to restrict action potential (AP) discharge, such as pattern separation (Sahay et al. 2011a; Sahay et al. 2011b). Indeed suppressing adult neurogenesis in mice appears to weaken pattern separation (Clelland et al. 2009; Nakashiba et al. 2012; Niibori et al. 2012; Tronel et al. 2012) and increasing adult neurogenesis improves it (Sahay et al. 2011a).

In addition, inhibition of the GC population by young adult-born GCs could limit excessive excitation from glutamatergic input and protect the cells in the DG hilus and hippocampus that are vulnerable to excitotoxicity. Thus, strong excitation of GCs can cause excitotoxicity of hilar neurons, area CA1 pyramidal cells, and area CA3 pyramidal cells (Scharfman and Schwartzkroin 1990b; a; Sloviter 1994; Scharfman 1999; Sloviter et al. 2003). Indeed, increasing adult-born neurons protects hilar neurons, and CA3 from neuronal loss 3 days after severe seizures are induced by the convulsant pilocarpine(Jain et al. 2019).

The seizures induced by kainic acid or pilocarpine are severe, continuous, and last several hours, a condition called *status epilepticus* (SE). The neuronal injury in hippocampus after SE has been suggested to be important because it is typically followed by chronic seizures (epilepsy) in rodents and humans, and has been suggested to cause the epilepsy (Falconer et al. 1964; Sloviter 1994; Cavalheiro et al. 1996; Herman 2002; Mathern et al. 2008; Dudek and Staley 2012; Dingledine et al.

2014). Chronic seizures involve the temporal lobe, so the type of epilepsy is called temporal lobe epilepsy (TLE). In the current study we asked if increasing adult -born neurons can protect from chronic seizures in an animal model of TLE. We used a very common method to induce a TLE-like syndrome, which involves injection of the muscarinic cholinergic agonist pilocarpine at a dose that elicits SE. Several weeks later, spontaneous intermittent seizures begin and continue for the lifespan (Scorza et al. 2009; Botterill et al. 2019; Levesque et al. 2021; Whitebirch et al. 2022). Seizure frequency, duration, and severity were measured by continuous video-EEG with 4 electrodes to monitor the hippocampus and cortex bilaterally.

It is known that SE increases adult neurogenesis (Parent and Kron 2012). SE triggers a proliferation of progenitors in the week after SE (Parent et al. 1997). Although many GCs that are born in the days after SE die in subsequent weeks by apoptosis, some survive. Young neurons that arise after SE and migrate into the GCL may suppress seizures by supporting inhibition of GCs because adult-born GCs in the normal brain inhibit GCs when they are young (Drew et al. 2016; Ash et al. 2023). In addition, after SE, the newborn GCs in the GCL can exhibit low excitability (Jakubs et al. 2006). However, some neurons born after SE mismigrate to ectopic locations such as the hilus (hilar ectopic GCs), where they can contribute to recurrent excitatory circuits that promote seizures (Scharfman et al. 2000; Parent and Lowenstein 2002; Scharfman 2004; Scharfman and Hen 2007; Parent and Murphy 2008; Scharfman and McCloskey 2009; Zhan et al. 2010; Myers et al. 2013; Cho et al. 2015; Althaus et al. 2019; Zhou et al. 2019). Since the hilar ectopic GCs are potential contributors to epileptogenesis, we also studied whether enhancing adult -born neurons would alter the number of hilar ectopic GCs.

Mossy cells are a major subset of glutamatergic hilar neurons which are vulnerable to excitotoxicity after SE (Scharfman 1999; Sloviter et al. 2003). During SE, mossy cells may contribute to the activity that ultimately leads to widespread neuronal loss (Botterill et al., 2019). However, surviving mossy cells can be beneficial after SE because they inhibit spontaneous chronic seizures in mice (Bui et al. 2018). Another large subset of vulnerable hilar neurons co-express GABA and somatostatin (SOM; Sloviter 1987; de Lanerolle et al. 1989; Freund et al. 1992; Sun et al. 2007) and correspond to so-called HIPP cells (neurons with ***<u>hi</u>***lar cell bodies and axons that project to the terminal zone of the ***<u>p</u>***erforant ***<u>p</u>***ath; (Han et al. 1993)). HIPP cells are important because they normally inhibit GCs and have the potential to prevent seizures. Therefore, we studied mossy cells and SOM cells in the current study.

The results showed that increasing adult -born neurons protects mossy cells and hilar SOM cells and reduces chronic seizures. Remarkably, the preservation of hilar mossy cells and SOM cells, and the reduction in chronic seizures, was found in females only. The sex difference may have been due to a greater ability to increase adult -born neurons in females than males, consistent with sex differences in *Bax*- and caspase-dependent cell death (Forger et al. 2004; Siegel and McCullough 2011).

The results are surprising because prior studies that suppressed neurogenesis reduces chronic seizures. Therefore, taken together with the results presented here, both increasing and suppressing adult-born neurons appear to reduce chronic seizures. How could this be? Past studies suggested that suppressing adult-born neurons led to a reduction in chronic seizures because there were fewer hilar ectopic granule cells. In the current study, increasing adult-born neurons may have reduced chronic seizures for another reason.Regardless, the present data suggest a novel and surprising series of findings which, taken together with past studies, suggest that adult--born neurons can be targeted in multiple ways to reduce chronic seizures in epilepsy.

## RESULTS

### I. Increasing adult-born neurons reduced the duration of pilocarpine-induced SE

#### **A.** General approach

The first experiment addressed the effect of increasing adult -born neurons on pilocarpine-induced SE in Nestin-CreER^T2^*Bax^fl/fl^*mice (called “Cre+”, below). To produce Nestin-CreER^T2^*Bax^fl/fl^* mice, hemizygous Nestin-CreER^T2^ mice were bred with homozygous *Bax^fl/fl^* mice. Littermates of Cre+ mice that lacked Cre (called “Cre-”, below) were also treated with tamoxifen and were controls.

Fig. 1A1 shows the experimental timeline. Tamoxifen was injected s.c. once per day for 5 days to delete *Bax* from Nestin-expressing progenitors. After 6 weeks, a time sufficient for a substantial increase in adult-born neurons (Drew et al., 2016, Jain et al., 2019), pilocarpine was injected s.c. to induce SE.

**Figure 1.**
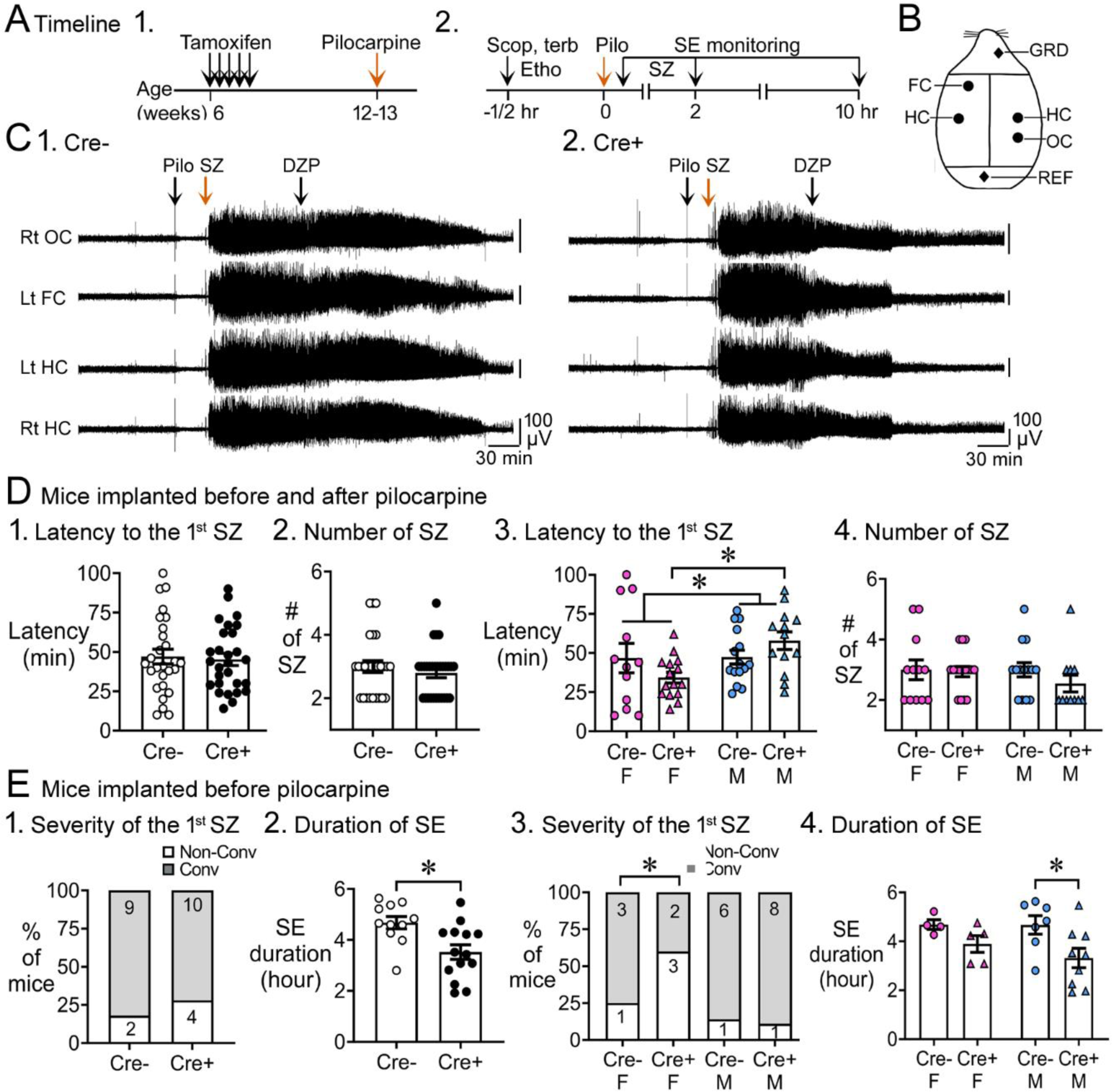
Pilocarpine-induced SE in Cre+ and Cre- mice. **A.** The experimental timeline is shown. 1. Tamoxifen was injected 1/day for 5 days in 6 week-old Nestin-CreER^T2^ *Bax^fl/fl^*mice. Six weeks after the last tamoxifen injection, mice were injected with pilocarpine (Pilo) at a dose that induces status epilepticus (SE). 2. On the day of pilocarpine injection, one group of mice without EEG electrodes were monitored for behavioral seizures for 2 hr after pilocarpine injection. Another group of mice were implanted with EEG electrodes 3 weeks prior to pilocarpine injection. In these mice, video-electroencephalogram (video-EEG) was used to monitor SE for 10 hr after pilocarpine injection. **B.** Locations to implant EEG electrodes are shown. Four circles represent recording sites: left frontal cortex (Lt FC), left hippocampus (Lt HC), right hippocampus (Rt HC) and right occipital cortex (Rt OC). Two diamonds represent ground (GRD) and reference (REF) electrodes. **D.** Pooled data for mice that were implanted with EEG electrodes and unimplanted mice. These data showed no significant genotypic differences but there was a sex difference. 1. The latency to the onset of first seizure was similar in both genotypes (t-test, p=0.761). The seizure was a behavioral seizure ≥stage 3 of the Racine scale (unilateral forelimb jerking). For this figure and all others, detailed statistics are in the Results. 2. The number of seizures in the first 2 hr after pilocarpine injection was similar in both genotypes (t-test, p=0.377). 3. After separating males and females, females showed a shorter latency to the onset of the first seizure compared to males (two-way ANOVA, p=0.043); Cre+ females had a shorter latency to the first seizure relative to Cre+ males (Bonferroni’s test, p=0.010). 4. The number of seizures in the first 2 hr after pilocarpine injection were similar in males and females (two-way ANOVA, p=0.436). **E.** Implanted mice. These data showed a significant protection of Cre+ mice on SE duration. 1. The severity of the first seizure (non-convulsive or convulsive) was similar between genotypes (Chi-square test, p=0.093). 2. Cre+ mice had a shorter duration of SE than Cre- mice (t-test, p=0.007). 3. After separating males and females, the first seizure was mostly non-convulsive in Cre+ females compared to Cre- females (60% vs. 14%) but no groups were statistically different (Fisher’s exact tests, p>0.05). 4. Once sexes were separated, there was no effect of sex by two-way ANOVA but a trend in Cre+ males to have a shorter SE duration than Cre- males (Bonferroni’s test, p=0.078).

Fig. 1A2 shows the experimental timeline during the day of pilocarpine injection. The location of electrodes for EEG are shown in Fig. 1B. Mice monitored with EEG were implanted with electrodes 3 weeks before SE (see Methods). Examples of the EEG are shown in Fig. 1C for Cre- and Cre+ mice and details are shown in Fig. 1- Supplemental figure 1.

#### **B.** Effects of increasing adult-born neurons on SE

The latency to the first seizure after pilocarpine injection was measured for all mice (with and without EEG electrodes; Fig. 1D) or just those that had EEG electrodes (Fig. 1E). When mice with and without electrodes were pooled, the latency to the onset of first seizure was similar in both genotypes (Cre-: 47.2 ± 4.8 min, n=27; Cre+: 45.3 ± 3.9 min, n=28; Student’s t-test, t(53)=0.3, p=0.761; Fig. 1D1).

The total number of seizures was quantified until 2 hr after pilocarpine injection because at that time diazepam was administered to decrease the severity SE. The total number of seizures were similar in both genotypes (Cre-: 3.0 ± 0.2 seizures; Cre+: 2.8 ± 0.1 seizures; Student’s t-test, t(54)=0.9, p=0.377; Fig. 1D2).

Interestingly, when sexes were separated, Cre+ females had a shorter latency to the first seizure than all other groups (Fig. 1D3). Thus, a two-way ANOVA with genotype (Cre- and Cre+) and sex (female and male) as main factors showed a main effect of sex (F(1,51)=4.31; p=0.043) with Cre+ females exhibiting a shorter latency compared to Cre+ males (Cre+ females, 34.3 ± 3.4 min, n=15; Cre+ males, 57.9 ± 5.6 min, n=13; Bonferroni’s test, p=0.026) but not other groups (Cre- female, 46.7 ± 9.3 min, n=12; Cre- males, 47.4 ± 4.4 min, n=15; Bonferroni’s tests, all p > 0.344; Fig. 1D3). There was no effect of genotype (F(1,49)=0.75; p=0.305) or sex (F(1,49)=0.62; p=0.436) on the total number of seizures by two-way ANOVA (Fig. 1D4).

When adult neurogenesis was suppressed by thymidine kinase activation in GFAP-expressing progenitors, the severity of the first seizure was worse, meaning it was often convulsive rather than non-convulsive (Iyengar et al., 2015). Therefore we examined the severity of the first seizure. These analyses were conducted only with mice implanted with electrodes because only with the EEG can one determine if a seizure is non-convulsive. A non-convulsive seizure was defined as an EEG seizure without movement. When sexes were pooled, the proportion of mice with a non-convulsive first seizure was not different (Cre-: 18.2%, 2/11 mice; Cre+: 28.6%, 4/14 mice Chi-square test, p>0.999; Fig. 1E1). However, when sexes were separated, the first seizure was non-convulsive in 60% of Cre+ females (3/5 mice) whereas only 25% of Cre- females had a first seizure that was nonconvulsive (1/4 mice), 14% of Cre- males (1/7 mice), and 11% of Cre+ males (1/9 mice; Fig. 1E3). Although the percentages suggest differences, i.e., Cre+ females were protected from an initial severe seizure, the differences were not significantly different (Fisher’s exact test, p=0.166; Fig. 1E3).

The duration of SE was shorter in Cre+ mice compared to Cre- mice (Cre-: 280.5 ± 14.6 min, n=11; Cre+: 211.4 ± 17.2 min, n=14; Student’s t-test, t(23)=0.30, p=0.007; Fig. 1E2). When sexes were separated, effects of genotype were modest. A two-way ANOVA showed that the duration of SE was significantly affected by genotype (F(1,21)=6.7; p=0.017) but not sex (F(1,21)=5.04; p=0.487). Cre+ males showed a trend for a shorter SE duration compared to Cre- males (Cre- males: 280.1 ± 22.8 min, n=7; Cre+ males: 199.1 ± 24.1 min, n=9; Bonferroni’s test, p=0.078; Fig. 1E4). Cre+ females had a mean SE duration that was shorter than Cre- females, but it was not a significant difference (Cre- females: 281.0 ± 11.8 min, n=4; Cre+ females: 233.4 ± 20.5 min, n=5; Bonferroni’s test, p=0.485; Fig. 1E4). More females would have been useful, but the incidence of SE in females was only 42.8% if they were implanted with EEG electrodes (Fig. 1 – Supplemental figure 2). In contrast, the incidence of SE in the unimplanted females was 100%, a significant difference by Fisher’s exact test (p < 0.001; Fig. 1- Supplemental figure 2). In males the incidence of SE was also significantly different in implanted and unimplanted mice (implanted males, 70.4%; unimplanted males, 100%; Fisher’s exact test, p=0.016, Fig. 1- Supplemental figure 2).

#### **D.** Power

We also investigated power during SE (Fig. 1- Supplemental figure 3). The baseline was measured, and then power was assessed for 5 hrs, at which time SE had ended. Power was assessed in 20 min consecutive bins. Females were used in this analysis (Cre- and Cre+). Two-way RMANOVA with genotype and time as main factors showed no effect of genotype for any frequency range: delta (1-4 Hz, F(1,7)=1.61; p=0.245); theta (4-8 Hz, F(1,7)=1.75; p=0.227); beta (8-30 Hz, F(1,7)=1.65; p=0.240); low gamma (80 Hz, F(1,7)=0.29; p=0.174); high gamma (80-100 Hz, F(1,7)=0.17; p=0.689). There was a significant effect of time for all bands (delta, p=0.003; theta, p=0.002, beta, low gamma and high gamma, p <0.001), which is consistent with the declining power in SE with time.

#### **E.** Role of diazepam

Diazepam was administered earlier in females during SE than males, and this could have influenced the results. However, the timing of SE was not significantly different in females than males (Fig. 1 – Supplemental figure 4). Also, diazepam was administered the same way in all Cre+ and Cre- females similarly but only the Cre+ females were protected as discussed below.

In summary, Cre+ mice did not show extensive differences in SE except for SE duration, which was shorter.

### II. Increasing adult-born neurons decreased chronic seizures

#### **A.** Numbers and frequency of chronic seizures

Continuous video-EEG was recorded for 3 weeks to capture chronic seizures (Fig. 2A). Representative examples of chronic seizures are presented in Fig. 2B. All chronic seizures were convulsive. First, we analyzed data with sexes pooled (Fig. 2C) and the total number of chronic seizures were similar in the two genotypes (Cre-: 22.6 ± 3.0 seizures, n=18; Cre+: 21.3 ± 1.6 seizures, n=17; Student’s t-test, t(33)=0.15, p=0.882; Fig. 2C1). The frequency of chronic seizures were also similar among genotypes (Cre-: 1.1 ± 0.14 seizures/day, n=18; Cre+: 1.0 ± 0.08 seizures/day, n=17; Welch’s t-test, t(26)=0.37, p=0.717; Fig. 2D1).

**Figure 2.**
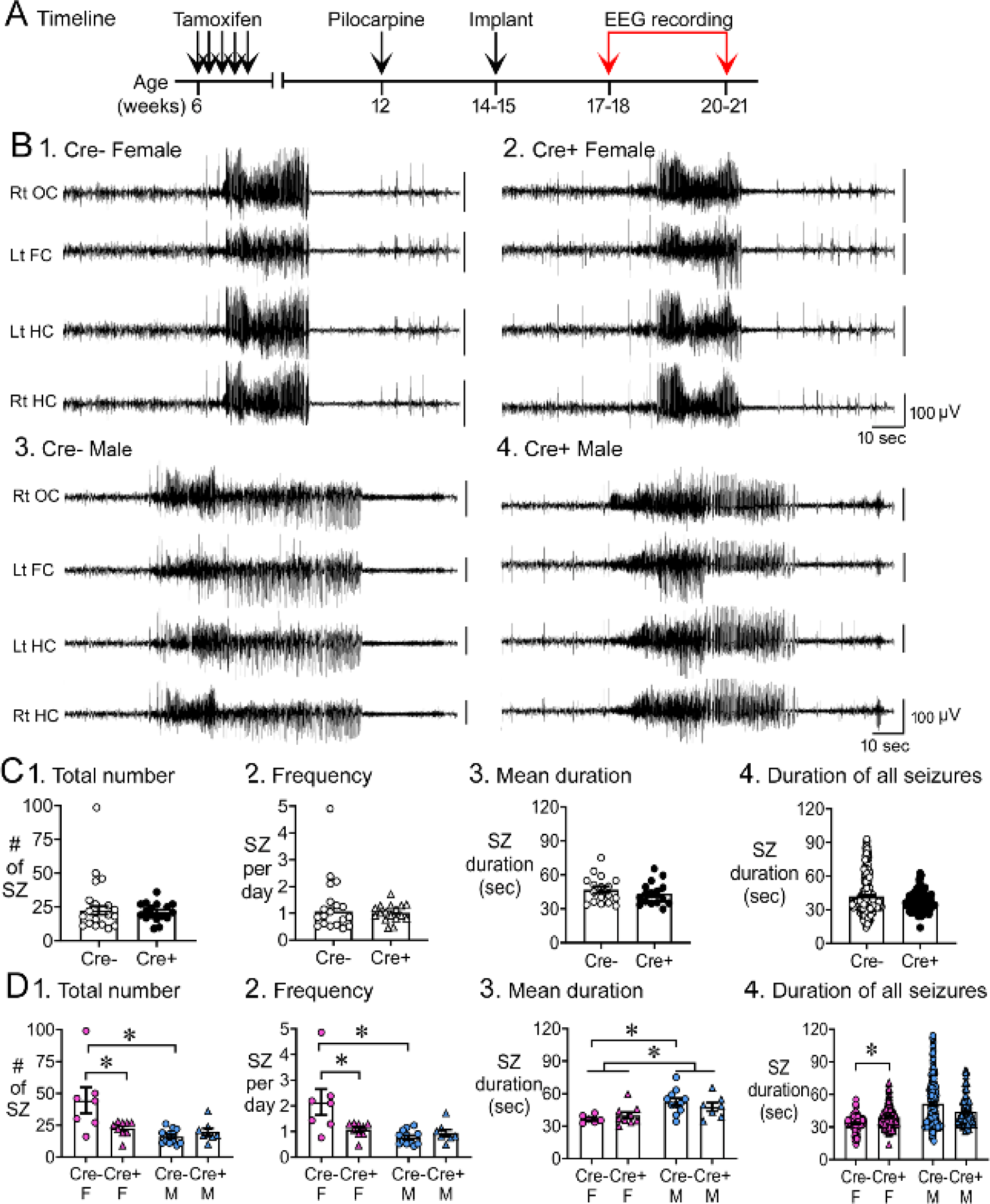
Reduced chronic seizures in Cre+ mice. A. The experimental timeline is shown. Six weeks after pilocarpine injection, continuous video-EEG was recorded for 3 weeks to capture chronic seizures. Mice that were unimplanted prior to SE were implanted at 2-3 weeks after pilocarpine injection. **B.** Representative examples of 2 min-long EEG segments show a seizure in a Cre- (1, 3) and Cre+ (2, 4) mouse. **C.** Numbers of chronic seizures. 1. Pooled data of females and males showed no significant effect of genotype on chronic seizure number. The total number of seizures during 3 weeks of recording were similar between genotypes (t-test, p=0. 882). 2. After separating data based on sex, females showed fewer seizures. Cre+ females had fewer seizures than Cre- females (Bonferroni’s test, p=0.004). There was a sex difference in control mice, with fewer seizures in Cre- males compared to Cre- females (Bonferroni’s test, p<0.001). **D.** Chronic seizure frequency. 1. Pooled data of females and males showed no significant effect of genotype on or chronic seizure frequency. The frequency of chronic seizures (number of seizures per day) were similar (Welch’s t-test, p=0.717). 2. Seizure frequency was reduced in Cre+ females compared to Cre- females (Bonferroni’s test, p=0.004). There was a sex difference in control mice, with lower seizure frequency in Cre- males compared to Cre- females (Bonferroni’s test, p<0.001). **E.** Seizure duration per mouse. 1. Each data point is the mean seizure duration for a mouse. Pooled data of females and males showed no significant effect of genotype on seizure duration (t-test, p=0.379). 2. There was a sex difference in seizure duration, with Cre- males having longer seizures than Cre- females (Bonferroni’s test, p=0.005). Because females exhibited more postictal depression (see Fig. 3), corresponding to spreading depolarization (Ssentongo et al. 2017), the shorter female seizures may have been due to truncation of seizures by spreading depolarization. **F.** Seizure durations for all seizures. 1. Every seizure is shown as a data point. The durations were similar for each genotype (Mann-Whitney *U* test, p=0.079). 2. Cre+ females showed longer seizures than Cre- females (Dunn’s test, p<0.001). Cre+ females may have had longer seizures because they were protected from spreading depolarization.

Data were then segregated based on sex and a two-way ANOVA was conducted with genotype and sex as main factors. There was a main effect of genotype (F(1,32)=4.26; p=0.047) and sex (F(1,32)=12.46; p=0.001) on the total number of chronic seizures and a significant interaction between sex and genotype (F(1,32)=8.54; p=0.006). Bonferroni’s post-hoc tests showed that Cre+ females had ∼half the chronic seizures of Cre- females (Cre- female: 44.6 ± 10.2 seizures, n=7; Cre+ female: 22.6 ± 2.0 seizures, n=9; p=0.004; Fig. 2C). However, Cre+ males and Cre- males had a similar number of chronic seizures (Cre- male: 16.1 ± 1.6 seizures, n=12; Cre+ male: 19.9 ± 2.7 seizures, n=8; p>0.999; Fig. 2C2).

Results for seizure frequency were similar to results comparing total numbers of seizures. There ws a main effect of genotype (F(1,32) 4.18; p=0.049) and sex (F(1, 32)=11.96; p=0.002) on chronic seizure frequency, and a significant interaction between sex and genotype (F(1,32)=8.29; p=0.007). Cre+ female mice had approximately half the seizures per day as Cre- females (Bonferroni’s test, Cre- female: 2.1 ± 0.5 seizures/day; Cre+ female: 1.1 ± 0.1 seizures/day; p=0.004; Fig. 2D2).

#### **B.** Additional analyses

While reviewing the data for each mouse plotted in Fig. 2C2 and 2D2, one point appeared spurious in the Cre- females, potentially influencing the comparison. The seizures in this mouse were more than 2x the standard deviation of the mean. Although not an outlier using the ROUT method (see Methods), we were curious if removing the data of this mouse would lead to a difference in the statistical results. There was still a main effect of sex (F(1,31)=16.04; p=0.0004) with a significant interaction between sex and genotype (F(1,31)=9.20; p=0.005) and Cre+ females had significantly fewer seizures than Cre- female mice (p=0.020; Fig. 2- Supplemental figure 1A1). Tests for seizure frequency led to the same conclusions (Fig. 2- Supplemental figure 1A2). These data suggest that spurious data point was not the reason for the results.

All mice were included in the analyses above, both those implanted and unimplanted during SE. Mice which were unimplanted prior to SE were implanted at approximately 2-3 weeks after pilocarpine to study chronic seizures. Because implantation affected the incidence of SE (discussed above), we asked if chronic seizures were different in implanted and unimplanted mice. The total number of chronic seizures (F(1,17)=1.33, p=0.265) and seizure frequency (F(1,17)=1.27, p=0.276) were similar, suggesting that implantation did not influence chronic seizures (Fig. 2- Supplemental figure 1B).

There were no significant differences in mortality associated with SE or chronic seizures. For quantification, we examined mortality during SE and the subsequent 3 days, 3 days until the end of the 3 week-long EEG recording period, or both (Fig. 2- Supplemental figure 2A). Graphs of mouse numbers (Fig. 2- Supplemental figure 2B) or percentages of mice (Fig. 2- Supplemental figure 2C) were similar: groups (Cre- females, Cre+ females, Cre- males, Cre+ males) were not significantly different (Chi-Sq. test, p>0.999).

#### **C.** Mean duration of individual chronic seizures

To evaluate the duration of individual seizures at the time mice were epileptic, two measurements were made. First, durations of each seizure of a given mouse were averaged, and then the averages for Cre- mice were compared to the averages for Cre+ mice (Fig. 2E1). There was no difference in the genotypes (Cre-: 46.8 ± 2.9 sec, n=17; Cre+: 43.4 ± 2.5 sec, n=16; Student’s t-test, t(31)=0.89, p=0.379; Fig. 2E1).

When separated by sex, a two-way ANOVA showed that female seizure durations were shorter than males (F(1,29)=12.42; p=0.001). However, this was a sex difference, not an effect of genotype (F(1,29)=0.033; p=0.856; Fig. 2E2), with Cre- female seizure duration shorter than Cre- male seizure duration (Bonferroni post-hoc test, p=0.015), and the same for Cre+ females compared to Cre+ males (Bonferroni post-hoc test, p=0.037; Fig. 2E2). One reason for the sex difference could be related to the greater incidence of postictal depression in females (see below), because that suggests spreading depolarizations truncated the seizures in females but not males.

The second method to compare seizure durations compared the duration of every seizure of every Cre- and Cre+ mouse. In the previous comparison (Fig. 2E1), every mouse was a data point, whereas here every seizure was a data point (Fig. 2F1). The data were similar between genotypes (Cre-: 41.9 ± 0.9 sec; Cre+: 36.8 ± 0.7 sec; Mann-Whitney *U* test, *U* statistic, 18873, p=0.079; Fig. 2F1). When sexes were separated, a Kruskal-Wallis test was significant (Kruskal-Wallis statistic, 69.30, p<0.001). Post-hoc tests showed that Cre+ females had longer seizure durations than Cre- females (Cre- female: 33.2 ± 0.7 sec; Cre+ female: 39.3 ± 0.6 sec, p<0.001; Fig. 2F2). Cre+ females may have had longer seizures because they were protected from spreading depolarizations that truncated seizures in Cre- females. Seizure durations were not significantly different in males (Cre- male: 51.4 ± 1.7 sec; Cre+ male: 44.0 ± 1.3 sec, p=0.298; Fig. 2F2).

#### **D.** Postictal depression

Postictal depression is a debilitating condition in humans where individuals suffer fatigue, confusion and cognitive impairment after a seizure. In the EEG, it is exhibited by a decrease in the EEG amplitude immediately after a seizure ends relative to baseline. In recent years the advent of DC amplifiers made it possible to show that postictal depression is often associated with spreading depolarization (Ssentongo et al. 2017), a large depolarization shift that is accompanied by depolarization block. As action potentials are blocked there are large decreases in input resistance leading to cessation of synaptic responses. As ion pumps are activated to restore equilibrium, there is recovery and the EEG returns to normal (Somjen 2001; Hartings et al. 2017; Herreras and Makarova 2020; Lu and Scharfman 2021).

We found that males had little evidence of postictal depression but it was common in females (Fig. 3), a sex difference that is consistent with greater spreading depolarization in females (Eikermann-Haerter et al. 2009; Bolay et al. 2011; Kudo et al. 2023). As shown in Fig. 3A1, a male showed a robust spontaneous seizure (selected from the 3 week-long recording period when mice are epileptic). However, the end of the seizure did not exhibit a decrease in the amplitude of the EEG relative to baseline. The EEG before and immediately after the seizure is expanded in Fig. 3A2 to show the EEG amplitude is similar. In contrast, the seizure from the female in Fig. 3B1-2 shows a large reduction in the EEG immediately after the seizure. For quantification, the mean peak-to-trough amplitude of the EEG 25-30 sec before the seizure was compared to the mean amplitude for the EEG during the maximal depression of the EEG after the seizure. If the depression was more than half, the animal was said to have had postictal depression.

**Figure 3.**
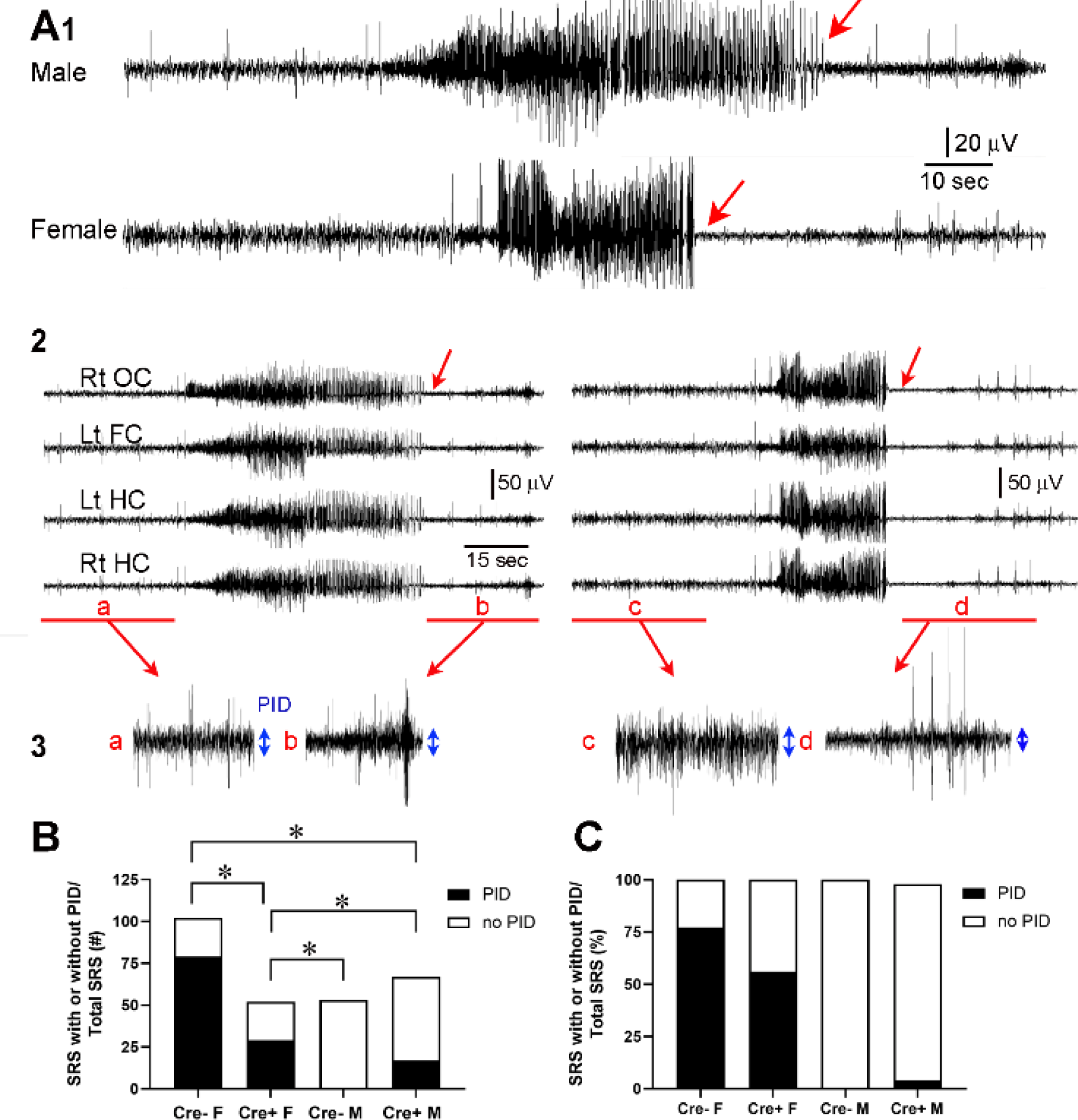
Reduced postictal depression in Cre+ female mice. A. 1. A seizure of a male mouse and female mouse are shown to illustrate postictal depression starting at the end of the seizure (red arrow). **2.** All 4 channels are shown for the male (left) and female mouse (right). Rt OC, right occipital cortex; Lt FC, left frontal cortex; Lt HC, left hippocampus; Rt HC, right hippocampus. The red arrows point to the end of the seizure. **3.** The areas in A2 marked by the red bar are expanded. The blue double -sided arrows reflect the mean EEG amplitude before (a, c) and after the seizure (b,d). **B.** For all spontaneous recurrent seizures (SRS) in the 3 week-long recording period, there was a significant difference between groups, with number of SRS with PID reduced in Cre+ females compared to Cre- females (Fisher’s exact test, all p <0.05). Males had very little postictal depression and there was no significant effect of genotype. **C.** The same data are plotted but the percentages are shown instead of the numbers of seizures.

When all chronic seizures were analyzed (n=274), the number of seizures with postictal depression was significantly different in the four groups (Cre- females, Cre+ females, Cre- males, Cre+ males; Chi-square test, p < 0.0001; Fig. 3C-D). Cre+ females showed less postictal depression compared to Cre- females (Fisher’s exact test, p=0.009; Fig. 3C). There was a sex difference, with females showing more postictal depression (108/154 seizures, 70.5%) than males (17/120 seizures, 14.2%; p < 0.0001; Fig. 3C-D).

#### **E.** Clusters of seizures

Next, we asked if the distribution of seizures during the 3 weeks of video-EEG was affected by increasing adult-born neurons. In Fig. 4A, a plot of the day-to-day variation in seizures is shown with each day of recording either black (if there were seizures) or white (if there were no seizures).

**Figure 4.**
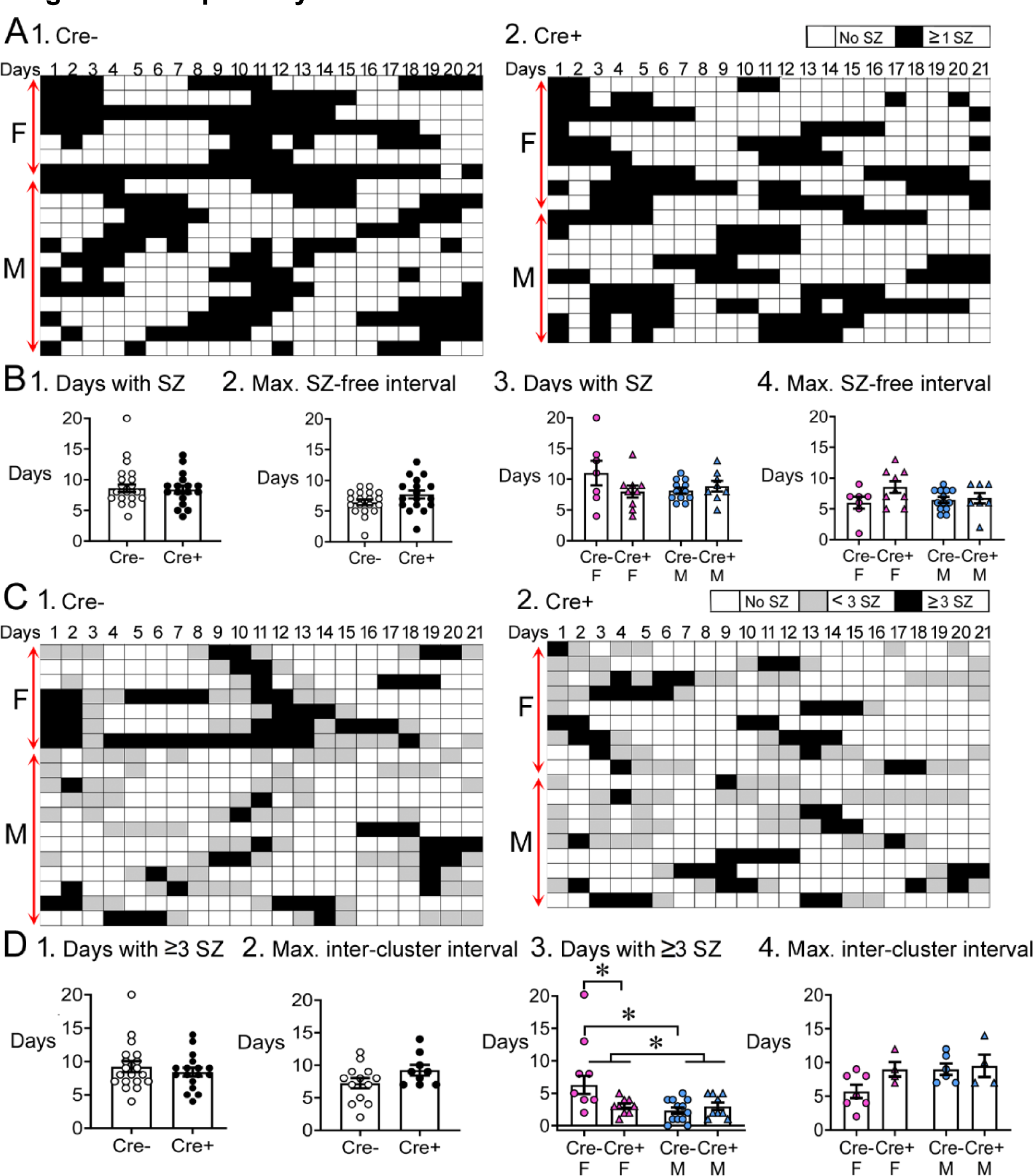
Temporal dynamics of chronic seizures. A. Each day of the 3 weeks-long EEG recording periods are shown. Each row is a different mouse. Days with seizures are coded as black boxes and days without seizures are white. 1. The number of days with seizures were similar between genotypes (t-test, p=0.822). 2. The maximum seizure-free interval was similar between genotypes (t-test, p=0.107). 3. After separating females and males, two-way ANOVA showed no effect of genotype or sex on days with seizures. 4. Two-way ANOVA showed no effect of genotype or sex on the maximum seizure- free interval. **C.** Then same data are shown but days with ≥3 seizures are black, days with < 3 seizures as grey, and are white. Clusters of seizures are reflected by the consecutive black boxes. 1. The cluster durations were similar between genotypes (Mann-Whitney’s *U* test, p=0.723). 2. The maximum inter-cluster interval was similar between genotypes (t-test, p=0.104. 3. Cre+ females had significantly fewer clusters than Cre- females (two-way ANOVA followed by Bonferroni’s test, p=0.009). There was a sex difference, with females having more clusters than males. Cre- females had more days with >3 seizures than control males (Cre- females: 6.3 ± 1.4 days; Cre- males: 2.3 ± 0.5 days; Bonferroni’s test, p < 0.001). 4. There was no significant effect of genotype or sex on the maximum inter-cluster interval. However, there was a trend for the inter-cluster interval to be longer in Cre+ females relative to than Cre- females.

The number of days with seizures were similar between genotypes (Cre-: 8.6 ± 0.6 days, n=18; Cre+: 8.4 ± 0.6 days, n=17; Student’s t-test, t(33)=0.23, p=0.822; Fig. 4B1). The number of consecutive days without seizures, called the seizure-free interval, was also similar between genotypes (Cre-: 6.4 ± 0.4 days, n=19; Cre+: 7.7 ± 0.6 days, n=17; Student’s t-test, t(34)=1.65, p=0.107; Fig. 4B2). When data were segregated based on sex, a two-way ANOVA showed no effect of genotype (F(1,32)=1.18, p=0.286) or sex on the number of days with seizures (F(1,32)=0.86, p=0.361; Fig. 4B3). There also was no effect of genotype (F(1,32)=2.86, p=0.100) or sex F(1,32)=0.53, p=0.471) on seizure-free interval (Fig. 4B4).

Clustering is commonly manifested by consecutive days with frequent seizures.

Clusters of seizures can have a substantial impact on the quality of life (Haut 2015; Jafarpour et al. 2019) so they are important. In humans, clusters are defined as at least 3 seizures within 24 hr (Goffin et al. 2007; Jafarpour et al. 2019). Therefore, we defined clusters as >1 consecutive day with ≥ 3 seizures/day (Fig. 4C). The duration of clusters were similar between genotypes (Cre-: 3.8 ± 0.7 days, n=19; Cre+: 3.0 ± 0.3 days, n=18; Mann-Whitney’s *U* test, *U* statistic 159, p=0.723; Fig. 4D1). Next, we calculated the number of days between clusters, which we call the intercluster interval. Genotypes were similar (Cre-: 7.2 ± 0.8 days, n=13; Cre+: 9.2 ± 0.8 days, n=9; Student’s t-test, t(20)=1.70, p=0.104; Fig. 4D2).

Two-way ANOVA was then performed on the sex-separated data. For cluster duration, there was no effect of genotype (F(1,33)=3.36, p=0.076) but there was a main effect of sex (F(1,33)=7.66, p=0.009) and a significant interaction of genotype and sex (F(1,33)=.66, p=0.009). Cre+ females had fewer days with ≥3 seizures than Cre- females (Cre- females: 6.3 ± 1.4 days; Cre+ females: 3.0 ± 0.4 days; Bonferroni’s test, p=0.009; Fig. 4D3). These data suggest Cre+ females were protected from the peak of a cluster, when seizures increase above 3/day.

There was no effect of genotype (F(1,17)=2.72, p=0.117) or sex (F(1,17)=2.72, p=0.117) on the intercluster interval (Fig. 4D4). However, this result may have underestimated effects because Cre+ females often had such a long interval that it was not captured in the 3 week-long recording period. That led to fewer Cre+ females that were included in the measurement of intercluster interval. In 5 out of 9 (i.e., 55%) Cre+ females, there was only one cluster in 3 weeks, so intercluster interval was too long to capture. Of those mice where intercluster interval could be measured, Cre- females had an interval of 5.7 ± 1.0 days (n=7) and Cre+ females had a 9.0 ± 1.1 day interval (n=4). That difference was not significant.

In summary, Cre+ females had fewer seizures, fewer days with ≥3 seizures, reduced postictal depression, and appeared to have a long period between clusters of seizures.

### III. Before and after epileptogenesis, Cre+ female mice exhibited more immature neurons than Cre- female mice but that was not true for male mice

#### **A.** Prior to SE

We first confirmed that prior to pilocarpine treatment, Cre+ mice had more young adult-born neurons compared to Cre- mice (Fig. 5A, Fig.5- Supplemental Fig. 1A-D). To that end, we quantified the adult-born GCs associated with the GCL/SGZ in both Cre+ and Cre- mice. DCX was used as a marker because it is highly expressed in immature neurons (Brown et al. 2003; Couillard-Despres et al. 2005). The area of the GCL/SGZ that exhibited DCX-ir was calculated and expressed as a percent of the total area of the GCL/SGZ (Fig. 5C).

**Figure 5.**
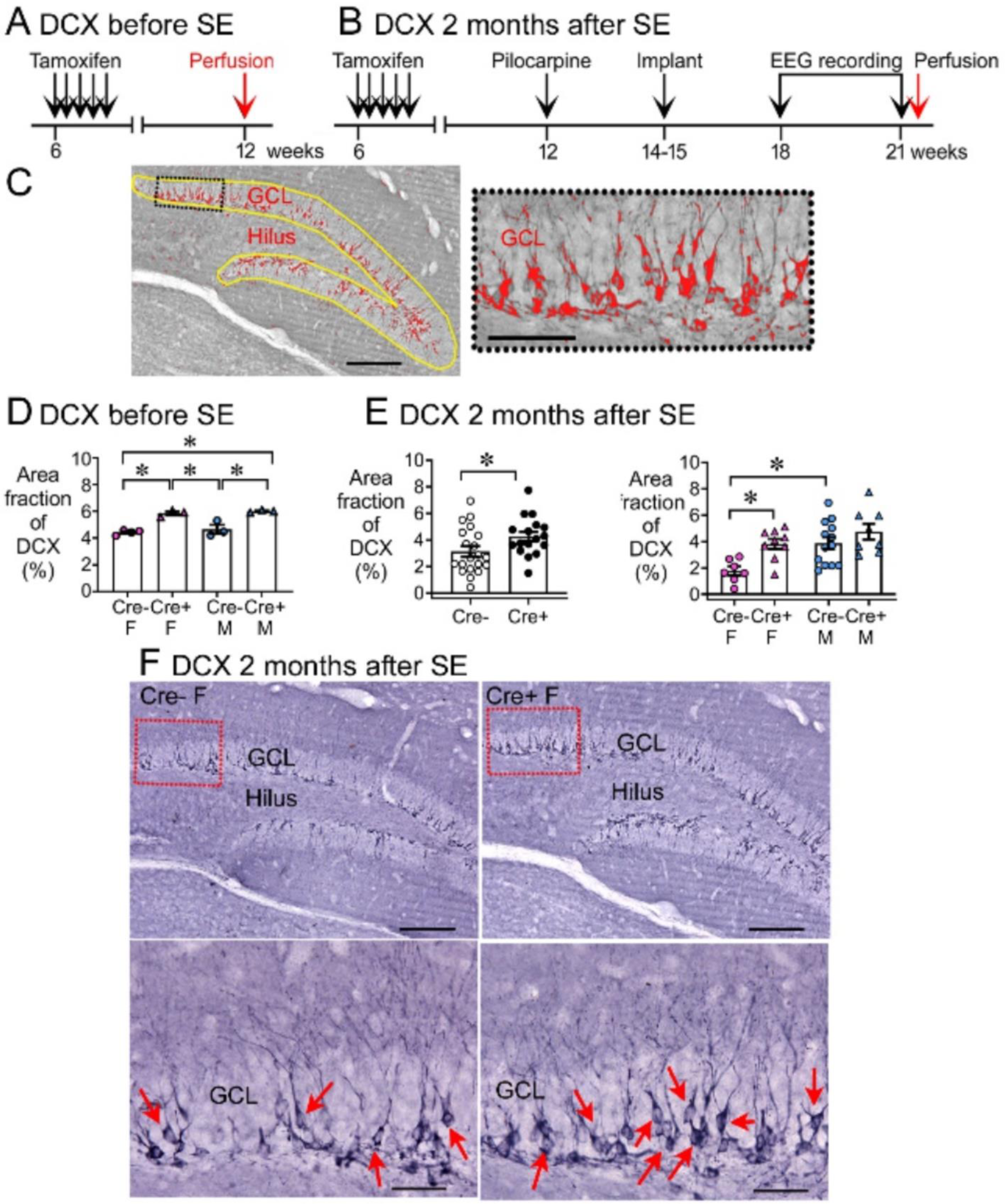
Increased DCX in Cre+ mice. A-B. The experimental timelines are shown. **A.** Mice were perfusion-fixed 6 weeks after tamoxifen injection, just before SE. Sections were then stained for DCX. **B.** Mice were tested 2 months after SE, after EEG recording. Then mice were perfused and staining was conducted for DCX. **B.** DCX quantification. DCX-ir within a region of interest (ROI; yellow lines) including the SGZ and GCL was thresholded. DCX-ir above the threshold is shown in red. Calibration, 100 μm (a); 50 μm (b). The inset is expanded to the right. **C.** The area of DCX-ir relative to the area of the ROI (referred to as area fraction) was greater in Cre+ mice compared to Cre- mice. Two-way ANOVA followed by Tukey pot-hoc tests, all p<0.05). **D.** Cre+ mice had increased DCX-ir relative to Cre- mice 2 months after SE. 1. Sexes were pooled. The area fraction of DCX-ir was greater in Cre+ than Cre- mice (t-test, p=0.041). 2. When sexes were separated, Cre+ females showed greater DCX-ir than Cre- females (two-way ANOVA followed by Bonferroni’s test, p=0.015). There was a sex difference, with Cre- males showing more DCX-ir than Cre- females (Bonferroni’s test, p=0.007). DCX-ir was similar in Cre- and Cre+ males (Bonferroni’s test, p=0.498). **E.** Representative examples of DCX-ir 2 months after SE. 1. Cre- female mouse. 2. Cre+ female mouse. The red boxes in a are expanded in b. Arrows point to DCX- ir cells. Calibration, 100 μm (a); 50 μm (b).

A two-way ANOVA with sex and genotype as factors showed a significant effect of genotype (F(1,9)=60.78, p<0.001) but not sex (F(1,9)=1.20, p=0.301; Fig. 5D). Post-hoc comparisons showed that Cre+ females had more DCX than Cre- females (p=0.001) and the same was true for males (p=0.003). Cre+ males had more DCX than Cre- females (p<0.001), and Cre+ females had more DCX than Cre- males (p=0.006). Cre- females and males were not different (p=0.774). The results are consistent with studies using the same methods which showed that Cre+ males have more DCX compared to Cre- males (Jain et al., 2019). Together the data suggest that Cre+ mice had more young adult-born neurons than Cre- mice immediately before SE.

#### **B.** After epileptogenesis

We also quantified DCX at the time when epilepsy had developed, after the 3 week-long EEG recording (Fig. 5B). Representative examples of DCX expression in the GCL/SGZ are presented in Fig. 5F and Fig. 5E shows the area fraction of DCX in the GCL/SGZ was significantly greater in Cre+ mice than Cre- mice (Cre-: 3.1 ± 0.4%, n=20; Cre+: 4.2 ± 0.3%, n=17; Student’s t-test, t(35)=2.13, p=0.041; Fig. 5D1).

Therefore, Cre+ mice had increased DCX in the GCL/SGZ after chronic seizures had developed.

To investigate a sex difference, a two-way ANOVA was conducted with genotype and sex as main factors. There was a significant effect of genotype (F(1,33)=12.62, p=0.001) and sex (F(1,33)=11.68, p=0.002), with Cre+ females having more DCX than Cre- females (Cre- female: 1.8 ± 0.3, n=7; Cre+ female: 3.8 ± 0.4, n=9; Bonferroni’s test, p=0.001; Fig. 5E). In contrast, DCX levels were similar between Cre+ and Cre- male mice (p=0.498, Fig. 5E). Therefore, elevated DCX occurred after chronic seizures had developed in Cre+ mice but the effect was limited to females. Because Cre+ epileptic females had increased immature neurons relative to Cre- females at the time of SE, and prior studies show that Cre+ females had less neuronal damage after SE (Jain et al. 2019), female Cre+ mice might have had reduced chronic seizures because of high numbers of immature neurons. However, the data do not prove a causal role.

It is notable that the Cre+ male mice did not show increased numbers of immature neurons at the time of chronic seizurers but Cre+ females did. It is possible that there was a “ceiling” effect in DCX expression that would explain why male Cre+ mice did not have a significant increase in immature neurons relative to male Cre- mice.

### IV. Hilar ectopic granule cells

Based on the literature showing that reducing hilar ectopic GCs decreases chronic seizures after pilocarpine-induced SE (Cho et al., 2015), we hypothesized that female Cre+ mice would have fewer hilar ectopic GCs than female Cre- mice. However, that female Cre+ mice did not have fewer hilar ectopic GCs.

To quantify hilar ectopic GCs we used Prox1 as a marker. Prox1 is a common marker of GCs in the GCL (Pleasure et al. 2000; Galeeva et al. 2007; Galichet et al. 2008; Steiner et al. 2008; Iwano et al. 2012), and the hilus (Scharfman et al. 2007; Hester and Danzer 2013; Cho et al. 2015; Bermudez-Hernandez et al. 2017).

Cre+ mice had significantly more hilar Prox1 cells than Cre- mice (Cre-: 19.6 ± 1.9 cells, n=18; Cre+: 60.5 ± 7.9 cells, n=18; Student’s t-test, t(34)=5.76, p<0.001; Fig. 6C1). A two-way ANOVA with genotype and sex as main factors showed no effect of sex (F(1,32)=0.28, p=0.595) but a significant effect of genotype (F(1,32)=0.23, p<0.0001) with more hilar Prox1 cells in female Cre+ than female Cre- mice (Cre- female: 18.2 ± 3.3 cells, n=7; Cre+ female: 57.0 ± 8.3 cells, n=9; Bonferroni’s test, p<0.009; Fig. 6C2) and the same for males (Cre- male: 20.4 ± 2.4 cells, n=11; Cre+ male: 63.9 ± 14.0 cells, n=9; Bonferroni’s test, p=0.001; Fig. 6C2).

**Figure 6.**
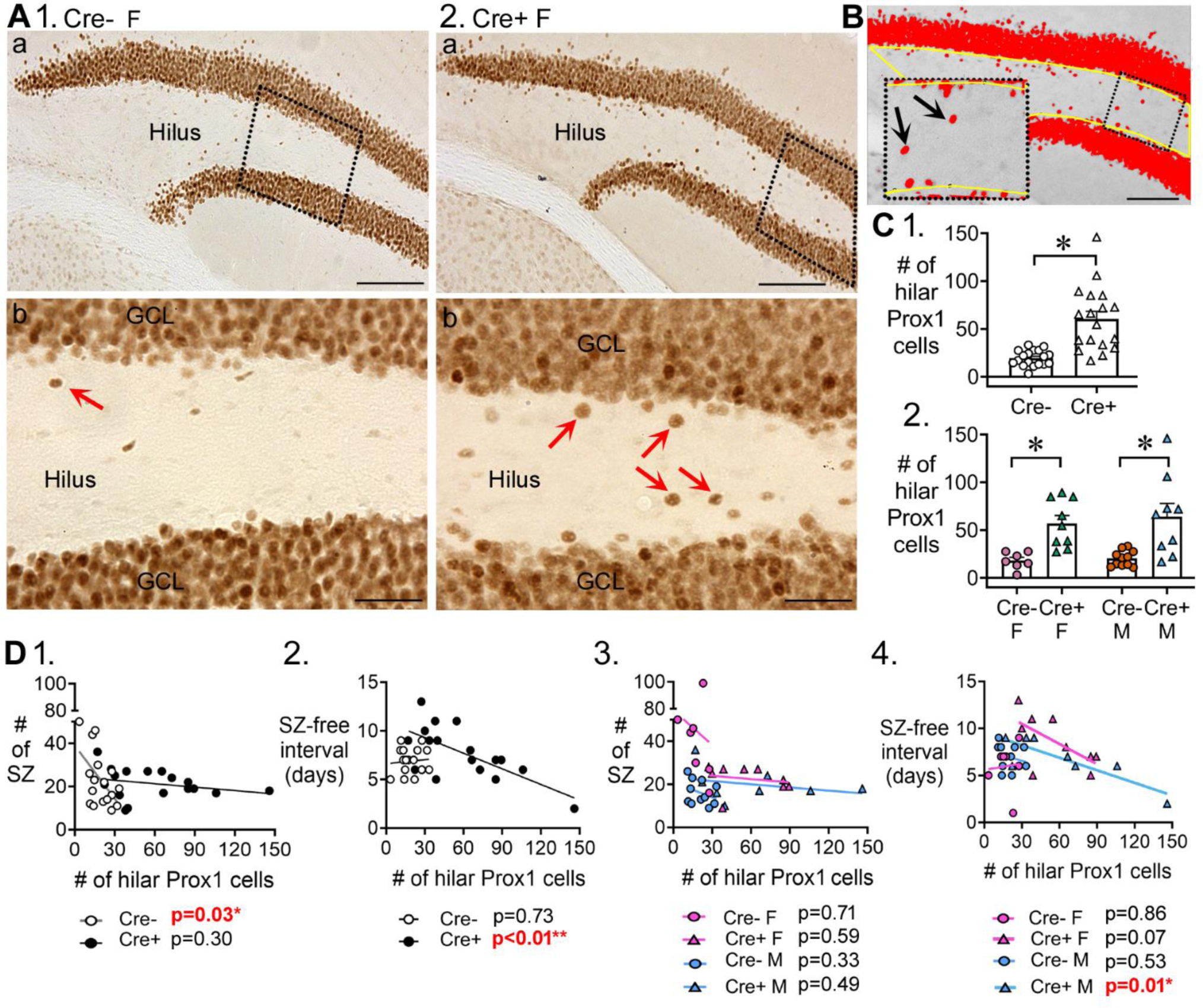
Hilar Prox1-ir cells increased in Cre+ mice. A. Representative examples of hilar Prox1-ir in Cre- (1) and Cre+ (2) mice are shown. The boxes in a are expanded in b. Arrows point to hilar Prox1-ir cells, corresponding to hilar ectopic GCs. Calibration, 100 μm (a); 50 μm (b). **B.** Prox1-ir is shown, within a hilar ROI. The area of the ROI above the threshold, relative to the area of the ROI, is red. This area is called the area fraction, and was used to quantify hilar Prox1-ir. Calibration, 100 μm. 1. Cre+ mice had more hilar Prox1-ir cells than Cre- mice (t-test, p<0.001). 2. When sexes were divided, Cre+ mice had more hilar Prox1-ir cells than Cre- mice in both female (two-way ANOVA followed by Bonferroni’s test, p<0.001) and male mice (Bonferroni’s test, p=0.001). **C.** Correlations between hilar Prox1-ir cells and measurements of chronic seizures. 1. All Cre- and Cre- mice were compared regardless of sex. For the Cre- mice there was a significant inverse correlation between the # of Prox1-ir cells and # of chronic seizures (R^2^=0.296). Thus, the more Prox1-ir cells there were, the fewer chronic seizures there were. However, that was not true for Cre+ mice (R^2^=0.072). 2. There was an inverse correlation between the number of hilar Prox1-ir cells and the seizure-free interval for Cre+ mice (R^2^=0.467) but not Cre- mice (R^2^=0.008). Thus, the more hilar Prox1-ir cells there were, the shorter the seizure-free periods were. However, this was not true for Cre- mice. 3. When data were divided by genotype and sex there was no significant correlation between hilar Prox1-ir cells and # of seizures (Cre- F, R^2^=0.0035; Cre+ F, R^2^=0.043; Cre- M, R^2^=0.104; Cre+ M, R^2^=0.083). 4. When data were divided by genotype and sex, there was a significant inverse correlation for the # of hilar Prox1-ir cells and seizure-free interval, but only for male Cre+ mice (R^2^=0.704). Cre+ females showed a trend (R^2^=0.395) and Cre- mice did not (Cre- F, R^2^=0.007, Cre- M, R^2^=0.046).

In past studies, hilar ectopic GCs have been suggested to promote seizures (Scharfman et al. 2000; Jung et al. 2006; Cho et al. 2015). Therefore, we asked if the numbers of hilar ectopic GCs correlated with the numbers of chronic seizures. When Cre- and Cre+ mice were compared (both sexes pooled), there was a correlation with numbers of chronic seizures (Fig. 6D1) but it suggested that more hilar ectopic GCs improved rather than worsened seizures. However, the correlation was only in Cre- mice, and when sexes were separated there was no correlation (Fig. 6D3).

When seizure-free interval was examined with sexes pooled, there was a correlation for Cre+ mice (Fig. 6D2) but not Cre- mice. Strangely, the correlations of Cre+ mice with seizure-free interval (Fig. 6D2, D4) suggest ectopic GCs shorten the seizure-free interval and therefore worsen epilepsy, opposite of the correlative data for numbers of chronic seizures. In light of these inconsistent results it seems that hilar ectopic granule cells had no consistent effect on chronic seizures.

### V. Increased adult-born neurons preserves mossy cells and hilar SOM interneurons but has little effect on parvalbumin interneurons

It has been suggested that epileptogenesis after a brain insult like SE is due to the hippocampal damage caused by the insult (Cavalheiro et al. 1996; Herman 2002; Mathern et al. 2008; Dudek and Staley 2012; Dingledine et al. 2014). Therefore, one of the reasons why increasing adult -born neurons reduced chronic seizures could be that it reduced neuronal damage after SE. Indeed, that has been shown (Jain et al., 2019). Here we examined the loss of vulnerable hilar mossy cells and SOM cells because they have been suggested to be critical (Sloviter 1987; Cavazos and Sutula 1990; Cavazos et al. 1994; Henshall and Meldrum 2012; Huusko et al. 2015). We asked whether Cre+ mice had preserved mossy cells (Fig. 7A) and SOM neurons (Fig. 7B). For comparison, we quantified the relatively seizure-resistant parvalbumin-expressing GABAergic neurons (Fig. 7C). An antibody to GluR2/3 was used as a marker of mossy cells (Leranth et al. 1996) and a SOM antibody for SOM cells (Leranth et al. 1990; Savanthrapadian et al. 2014; Botterill et al. 2019).

**Figure 7.**
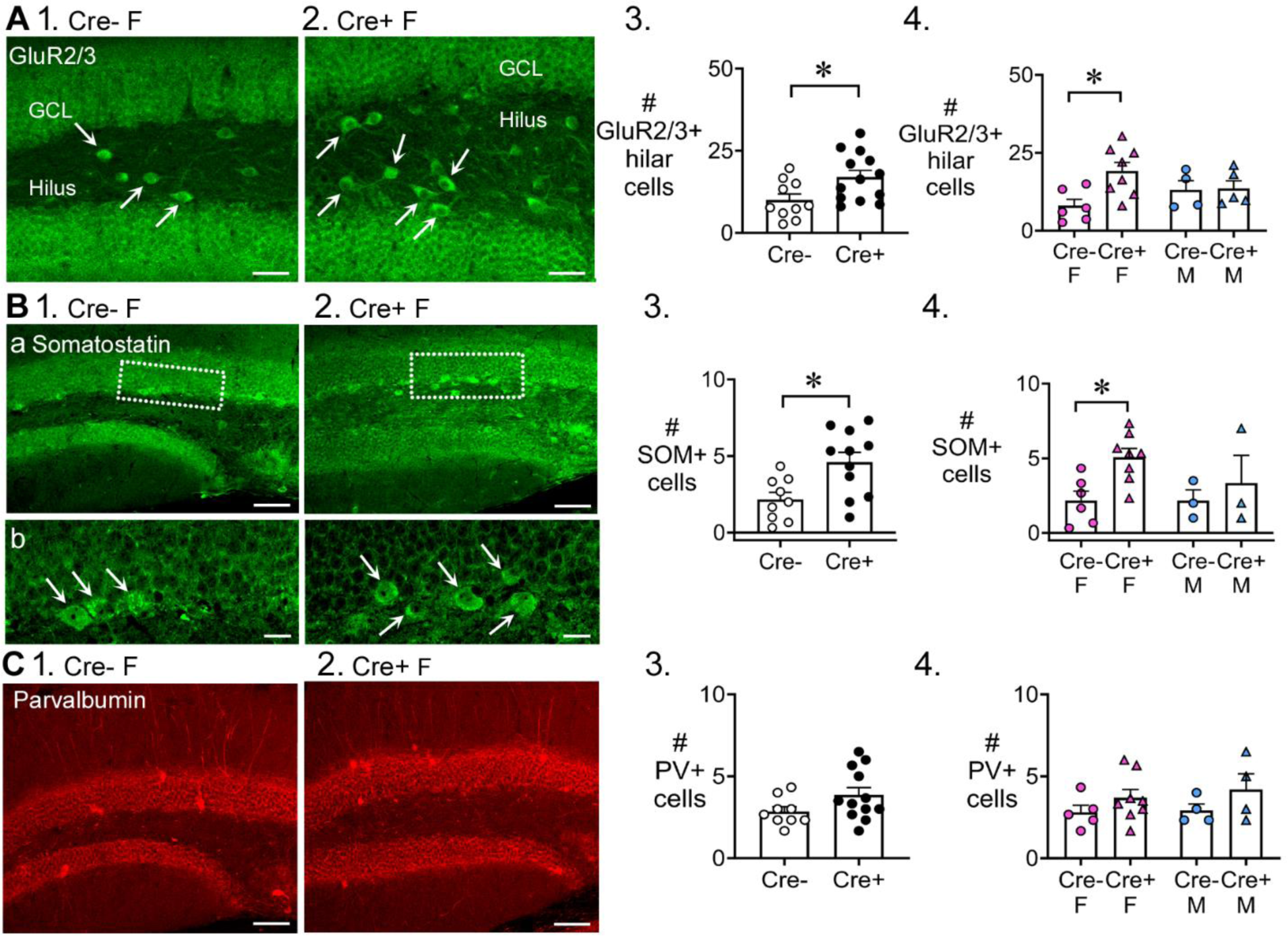
Preserved mossy cells and hilar SOM cells in Cre+ female mice but not parvalbumin interneurons. **A.**1-2. Representative examples of GluR2/3 labelling of Cre- (1) and Cre+ mice (2). Calibration, 50 μm. 3. Cre+ mice had more hilar GluR2/3-immunofluorescent (positive; +) cells than Cre- mice (t-test, p=0.022). Sexes were pooled. 4. After separating females and males, Cre+ females showed more hilar GluR2/3+ cells than Cre- females (Bonferroni’s test, p=0.011). Hilar GluR2/3+ cells were similar between genotypes in males (Bonferroni’s test, p=0.915). **B.** 1-2. Representative examples of SOM labelling in Cre- and Cre+ mice are shown. Calibration, 100 μm (a); 20 μm (b). 3. In pooled data, Cre+ mice had more hilar SOM cells than Cre- mice (t-test, p=0.008). 4. After separating females and males, Cre+ females showed more hilar SOM cells than Cre- females (Bonferroni’s test, p=0.019). Hilar SOM cells were similar between genotypes in males (Bonferroni’s test, p=0.897). **C.** 1-2. Representative examples of parvalbumin labelling in Cre- and Cre+ mice are shown. Calibration, 100 μm. 3. The number of parvalbumin+ cells in the DG were similar in Cre- and Cre+ mice in pooled data (t-test, p=0.095). 4. There was no effect of genotype (p=0.096) or sex (p=0.616) on the number of DG parvalbumin+ cells.

The results showed that Cre+ mice had more GluR2/3-expressing hilar cells than Cre- mice (Cre-: 10.0 ± 1.8 cells, n=10; Cre+: 17.0 ± 2.0 cells, n=13; Student’s t-test, t(21)=2.46, p=0.022; Fig. 7A1-3). We confirmed that the GluR2/3+ hilar cells were not double-labeled with Prox1, suggesting they corresponded to mossy cells, not hilar ectopic GCs (Supplementary Fig. 8A). To investigate sex differences, a two-way ANOVA was conducted with genotype and sex as main factors. There was a significant effect of genotype (F(1,18)=4.95, p=0.039) with Cre+ females having more GluR2/3 cells than Cre- females (Cre- female: 8.0 ± 2.0 cells, n=6; Cre+ female: 19.1 ± 2.7 cells, n=8; Bonferroni’s test, p=0.011; Fig. 7A4). GluR2/3-ir hilar cells were similar in males (Cre- male: 13.0 ± 3.0 cells, n=4; Cre+ male: 13.6 ± 2.5 cells, n=5; Bonferroni’s test, p=0.915; Fig. 7A4). These results in dorsal DG also were obtained in ventral DG (Supplementary Fig. 8B-C). The data suggest that having more GluR2/3-ir mossy cells could be a mechanism that allowed Cre+ females to have reduced chronic seizures compared to Cre- females. Equal numbers of GluR2/3 mossy cells in Cre+ and Cre- males could relate to their lack of protection against chronic seizures.

Next, we measured SOM hilar cells in pooled data (females and males together).

These results were analogous to the data for GluR2/3, showing that Cre+ mice had more hilar SOM cells than Cre- mice (Cre-: 2.1 ± 0.5 cells, n=9; Cre+: 4.6 ± 0.6 cells, n=11; Student’s t-test, t(18)=2.95, p=0.008; Fig. 7B1-3). When sexes were separated, a two-way ANOVA showed a significant effect of genotype (F(1,16)=5.14, p=0.038) and no effect of sex (F(1,18)=0.94, p=0.346). However, Cre+ females had more SOM cells than Cre- females (Cre- female: 2.2 ± 0.6 cells, n=6; Cre+ female: 5.1 ± 0.6 cells, n=8; Bonferroni’s test, p=0.019; Fig. 7B4), although only in dorsal DG (Fig. 7B4) not ventral DG (Supplementary Fig. 8C). Numbers of SOM cells were similar in males (Cre- male: 2.2 ± 0.7 cells, n=3; Cre+ male: 3.3 ± 1.8 cells, n=3; Bonferroni’s test, p=0.897; Fig. 7B4) in both dorsal and ventral DG (Supplementary Fig. 8B-C). Therefore, the ability to preserve more mossy cells and SOM hilar cells in Cre+ females could be a mechanism by which Cre+ females were protected from chronic seizures.

Parvalbumin-ir cells were not significantly different between genotypes (Student’s test, t(19)=1.76, p=0.095; Fig. 7C1-3). A two-way ANOVA showed no effect of genotype (F(1,17)=3.10, p=0.096) or sex (F(1,17)=0.26, p=0.616) on the numbers of parvalbumin cells. The results were the same in dorsal and ventral DG (Supplementary Fig. 8B-C).

These data are consistent with the idea that loss of parvalbumin-expressing cells has not been considered to play a substantial in epileptogenesis in the past (Sloviter 1987; 1994). However, it should be noted that subsequent research has shown that the topic is complicated because parvalbumin expression may decline even if the cells do not die (Andre et al. 2001; Sun et al. 2007) and data vary depending on the animal model (van Vliet et al. 2004; Huusko et al. 2015).

### VI. Increased adult-born neurons decreased neuronal damage after SE

In our previous study of Cre+ and Cre- mice (Jain et al., 2019), tamoxifen was administered at 6 weeks and SE was induced at 12 weeks (like the current study). We examined neuronal loss 3 days after SE, when neuronal loss in the hilus and area CA3 is robust in wild type mice. There is also some neuronal loss in CA1 at 3 days but more at 10 days after SE. We found less neuronal loss in Cre+ mice in these three areas (Jain et al. 2019). In the current study we examined 10 days after SE (Fig. 8A-B) because at this time delayed neuronal loss occurs, providing a better understanding of CA1 and the subiculum because delayed cell death occurs there. The intent was to determine if Cre+ mice exhibited less neuronal loss or not in CA1 and the subiculum.

**Figure. 8.**
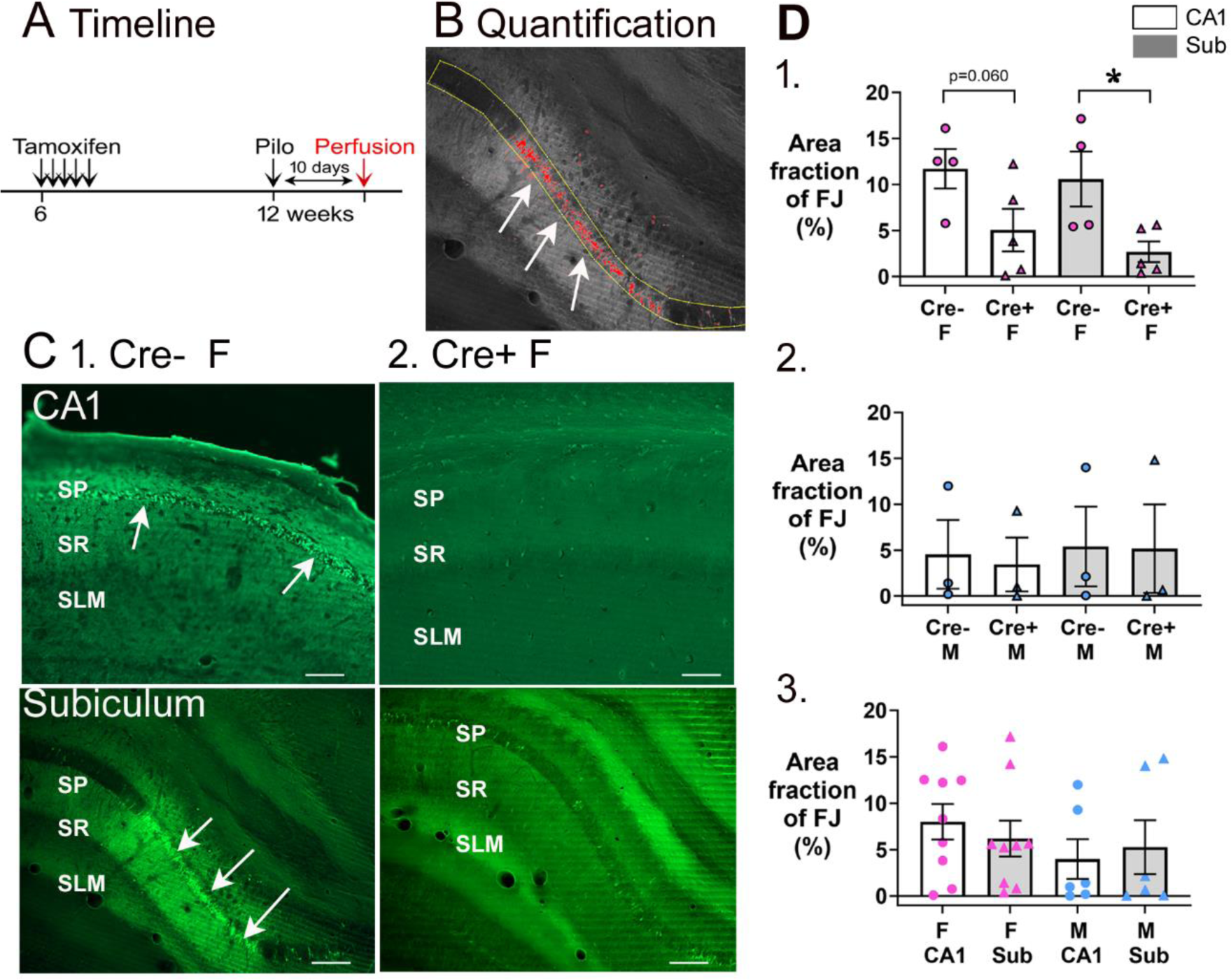
Cre+ female mice had less neuronal loss in hippocampus after SE. A. A timeline is shown to illustrate when mice were perfused to examine Fluorojade- C staining. All mice were perfused 10 days after SE, a time when delayed cell death occurs after SE, mainly in area CA1 and subiculum. Note that prior studies showed hilar and CA3 neurons, which exhibit more rapid cell death after SE, are protected from cell loss in Cre+ mice examined 3 days after SE (Jain et al., 2019). Also, there was protection of CA1 at 3 days (Jain et al., 2019). **B.** Quantification. Fluorojade-C was thresholded using ImageJ and the pyramidal cell layer outlined in yellow. The fraction above threshold relative to the entire ROI (area fraction) was calculated (see Methods) 2. The Fluorojade-C area fraction was greater in Cre- mice than Cre+ mice. Statistical comparisons showed a trend for CA1 of Cre- mice to exhibit more Fluorojade-C than Cre+ mice (Mann-Whitney *U* test, p=0.060). Cre- mice had a significantly greater area fraction in the subiculum than Cre+ mice (Mann- Whitney *U* test, p=0.032). **C.** Examples of Fluorojade-C staining in CA1 (top) and subiculum (bottom) of Cre+ female (1) and Cre- female (2) mice. SO, stratum oriens; SP, stratum pyramidale; SR, stratum radiatum; SLM, stratum lacunosum-moleculare. Arrows point to numerous Fluorojade-C-stained neurons in Cre- mice but not Cre+ mice. Calibration, 200 μm. **D.** 1. Comparisons of female mice by two-way ANOVA showed an effect of genotype (F(1,15)=11.97, p=0.004) with less Fluorojade C in Cre+ mice for CA1 (p=0.016) and subiculum p=0.016). 2. Comparisons of male mice showed no significant effect of genotype on Fluorojade C in either CA1 or the subiculum (F(1,8)=0.002, p=0.965; CA1, p=0.828, subiculum, p=0.973, respectively). 3. When genotypes were pooled, female mice did not have significantly more damage than males (two-way ANOVA, sex (F(1,34)=3.16, p=0.085) and there was no effect of subfield (F(1,34)=0.0016, p=0.968).

To quantify Fluorojade-C, ROIs were drawn digitally around the pyramidal cell layers (Fig. 8B). As shown in Fig. 8C, there was less Fluorojade-C staining in Cre+ female mice relative to Cre- female mice in both CA1 and the subiculum. The area of the ROI that showed Fluorojade C-positive cells was calculated as area fraction and expressed as % in Fig. 8D. For females, a two-way ANOVA with genotype and subfield as factors showed a significant effect of genotype (F(1,14)=11.21, p=0.005) with a smaller area fraction in Cre+ mice than Cre- mice for subiculum (p=0.045) and but not CA1 (p=0.095; Fig. 8D1). Males showed no significant differences either in genotype (F(1,8)=0.06, p=0.816) or subfield (F(1,8)=0.05, p=0.825; Fig. 8D2). When genotypes were pooled, females were not different than males either for CA1 or the subiculum (two-way ANOVA, sex, F(1,37)=0.466, p=0.499; Fig. 8D3). Thus, Cre+ female mice were protected from hilar, CA3, and CA1 damage at 3 days and were protected from subicular damage at 10 days after SE.

## DISCUSSION

This study showed that conditional deletion of *Bax* from Nestin-expressing progenitors increased young adult-born neurons in the DG when studied 6 weeks after deletion and using DCX as a marker of immature neurons. In a different set of mice, pilocarpine was used to induce epileptogenesis. The chronic seizures, measured 4-7 weeks after pilocarpine, were reduced in frequency by about 50% in females. Therefore, increasing young adult-born neurons before the epileptogenic insult can protect against epilepsy. However, we do not know if the protective effect was due to the greater number of new neurons before SE or other effects. Past data would suggest that increased numbers of newborn neurons before SE leads to a reduced SE duration and less neuronal damage in the days after SE. That would be likely to lessen the epilepsy after SE. However, there may have been additional effects of larger numbers of newborn neurons prior to SE.

Increasing young adult-born neurons has been shown to protect the hippocampus from SE-induced neuronal loss (Jain et al., 2019) which is a major contributor to epileptogenesis. Therefore, by protecting against SE-induced neuronal loss the young adult-born neurons could have reduced the severity of epilepsy. Indeed, we showed that the Cre+ female mice that had reduced chronic seizures had preservation of hilar mossy cells and SOM cells, two populations that are lost in SE- induced epilepsy and considered to contribute to epileptogenesis.

There were major surprises in the current study. First, the results are unexpected because suppressing adult-born neurons was shown to reduce chronic seizures (Cho et al., 2015). Here, increasing adult-born neurons did not have the opposite effect.

It was also unanticipated that only females with *Bax* deletion showed a significant increase in young adult-born neurons and a significant reduction in chronic seizures. The larger effect of increasing adult-born neurons in female mice may be attributable to sex differences in Bax (discussed below). The other remarkable finding relates to hilar ectopic GCs. These GCs have been suggested to promote epilepsy (Scharfman 2004; Jung et al. 2006; Scharfman et al. 2007; Parent and Murphy 2008; Hester and Danzer 2013; Cho et al. 2015), but hilar ectopic GCs increased in females with reduced seizures. The association of more hilar ectopic GCs with fewer chronic seizures was unexpected.

### Effects of *Bax* deletion on SE

In past studies, suppressing adult-born neurons made kainic acid-induced SE worse, and pilocarpine-induced SE was also worse (Iyengar et al., 2015; Jain et al., 2019). In the present study, SE was affected also. The duration of SE was reduced in Cre+ mice. In the Cre+ females, the first seizure after pilocarpine injection was often less severe, and power showed a tendency to be reduced during SE. Therefore, SE might have been less severe in the Cre+ females, and this could have contributed to reduced neuronal loss and chronic seizures.

### Chronic seizures

It is remarkable that increasing adult-born neurons for 6 weeks was sufficient to reduce seizures long-term. It is consistent with the idea that normally the young adult- born neurons inhibit other GCs, which supports the DG gate function (Hsu 2007; Drew et al. 2016). This gate has been suggested to be an inhibitory barrier to entry of seizures from cortex into hippocampus (Coulter and Carlson 2007; Hsu 2007; Krook- Magnuson et al. 2015).That entry is deleterious because seizures that pass from entorhinal cortex to the GCs and then CA3 are likely to continue to CA1 and back to cortex, causing reverberatory (long-lasting, severe) seizures. The reason for the relatively ease of reverberation once past the DG gate is that the synapses between GCs and CA3, CA1 and cortex are excitatory. The GCs have especially powerful excitatory synapses on CA3 pyramidal cells (Henze et al. 2000; Scharfman and MacLusky 2014), although these are normally mitigated by GABAergic circuitry (Acsady et al. 1998).

These data are consistent with the demonstration that adult-born neurons protect against other pathological conditions such as Alzheimer’s disease (Choi et al. 2018; Choi and Tanzi 2019). However, it is important to note that all effects are unlikely to be mediated only by the DG. The olfactory bulb and other areas also have adult-born neurons and they could contribute to epilepsy, especially those epilepsy syndromes with mechanisms that are extrahippocampal.

### Clusters of seizures

There were fewer days with > 3 seizures in Cre+ female mice which is another way that Cre+ females were protected from chronic seizures. These findings are valuable because clusters in humans have a significantly negative impact on health and quality of life (Haut 2015; Jafarpour et al. 2019).

The results may have underestimated the effects on clusters because we did not measure the interval between clusters in many Cre+ female mice. The reason is that the interval between clusters increased in some mice so they only had one cluster in 3 weeks. Thus intercluster interval appeared to lengthen in Cre+ females but animals with only one cluster had to be excluded. In the end, the results were not statistically significant.

### Sex differences

Females showed more of an effect of conditional *Bax* deletion than males. Insight into this sex difference came when the same assessments were made before SE Before SE, there was no sex difference. Cre+ females had more adult-born neurons than Cre- females and Cre+ males had more than Cre- males. In addition, the levels of DCX were similar in Cre+ females and Cre+ males.

However, after epilepsy developed, there was a sex difference. Cre- females had less DCX than Cre- males. One explanation is that cell birth during epileptogenesis was greater in males because it is in developing hippocampus (Sisk et al. 2016) and SE has been suggested to rekindle developmental programs (Ben-Ari and Holmes 2006).

Another possibility is males had less programmed cell death during epileptogenesis. Indeed during development, females have more apoptotic profiles than males and the sex difference was blocked by *Bax* deletion (Forger et al. 2004). A final possibililty is that cell death during epileptogenesis is *Bax*-dependent in females but *Bax*-independent in males. Support for this idea comes from studies of ischic cell death, which is caspase-dependent cell death in females but not males (Siegel and McCullough 2011).

### Hilar ectopic GCs

In the normal brain, adult-born neurons in the DG are thought to arise mainly from the SGZ and migrate to the GCL(Kempermann 2012). After SE, there is a surge in proliferation in the SGZ and neurons either migrate correctly to the GCL or aberrantly in the hilus (Parent et al. 1997).

These hilar ectopic GCs are thought to contribute to seizure generation in the epileptic brain because they are innervated by residual CA3 neurons, and project to GCs, making a major contribution to mossy fiber innervation of GCs in the inner molecular layer (Scharfman et al. 2000; Kron et al. 2010; Pierce et al. 2011; Scharfman and Pierce 2012; Althaus et al. 2016). When epileptiform activity occurs in CA3 in slices of epileptic rats, CA3 evokes discharges in hilar ectopic GCs that in turn excite GCs in the GCL (Scharfman et al. 2000). Consistent with the idea that hilar ectopic GCs promote seizures, the numbers of hilar ectopic GCs are correlated with chronic seizure frequency in rats (McCloskey et al. 2006) and mice (Hester and Danzer 2013).

Furthermore, suppressing hilar ectopic GC formation reduces chronic seizures (Jung et al. 2006; Cho et al. 2015; Hosford et al. 2016).

Notably, this is the first study to our knowledge showing that increased hilar ectopic GCs were found in mice that had reduced seizures. One potential explanation is that SE-induced hippocampal damage was reduced in Cre+ females with high numbers of hilar ectopic GCs. Therefore, the circuitry of the DG would be very different compared to past studies of hilar ectopic GCs where neuronal loss was severe (Supplementary Fig. 7). The presence of mossy cells is one way the circuitry would be different. MCs normally support the young adult-born GCs that migrate to the GCL (Piatti and Schinder 2018). Mossy cells provide an important activator of newborn GCs when they are young (Chancey et al. 2014). Mossy cells also innervate hilar ectopic GCs (Pierce et al. 2007).

Another possibility is that there was protection against chronic seizures in female Cre+ mice by increasing adult-born neurons *in the GCL*. The reason to suggest this possibility is that prior studies showed that young adult-born neurons in the GCL primarily inhibit GCs in the normal brain (Drew et al. 2016) and are relatively quiescent in the epileptic brain (Jakubs et al. 2006) .

### Additional considerations

This study is limited by the possibilities of type II statistical errors in those instances where we divided groups by genotype and sex, leading to comparisons of 3-5 mice/group. Another potential caveat is that female mice were selected regardless of the stage of the estrous cycle.

## Conclusions

In the past, suppressing adult neurogenesis before SE was followed by fewer hilar ectopic GCs and reduced chronic seizures. Here, we show that the opposite - enhancing adult-born neurons before SE and increased hilar ectopic GCs - do not necessarily reduce seizures. We suggest instead that protection of the hilar neurons from SE-induced excitotoxicity was critical to reducing seizures. The reason for the suggestion is that the survival of hilar neurons would lead to persistence of the normal inhibitory functions of hilar neurons, protecting against seizures. However, this is only a suggestion at the present time because we do not have data to prove it. Additionally, because protection was in females, sex differences are likely to have played an important role. Regardless, the results show that enhancing-born neurons of young adult-born neurons in Nestin-Cre+ mice had a striking effect in the pilocarpine model, reducing chronic seizures in female mice.

## MATERIALS AND METHODS

### I. General information

Animal care and use was approved by the Nathan Kline Institute Institutional Animal Care and Use Committee and met the regulations of the National Institute of Health and the New York State Department of Health. Mice were housed in standard mouse cages, with a 12 hr light/dark cycle and food (Laboratory rodent diet 5001; W.F. Fisher & Sons) and water *ad libitum*. During gestation and until weaning, mice were fed chow formulated for breeding (Formulab diet 5008; W.F. Fisher & Sons).

### II. Increasing adult-born neurons

To enhance-born neurons, a method was used that depends on deletion of *Bax*, the major regulator of programmed cell death in adult-born neurons (Sun et al. 2004; Sahay et al. 2011a; Ikrar et al. 2013; Adlaf et al. 2017). Enhancement of-born neurons was induced by conditional deletion of *Bax* from Nestin-expressing progenitors (Sahay et al. 2011a). These mice were created by crossing mice that have *loxP* sites flanking the pro-apoptotic gene *Bax* (*Bax^fl/fl^*) with a Nestin-CreER^T2^ mouse line in which tamoxifen-inducible Cre recombinase (CreER^T2^) is expressed under the control of the rat *Nestin* promoter (Sahay et al. 2011a). It was shown that after tamoxifen injection in adult mice there is an increase in dentate gyrus neurogenesis based on studies of bromo-deoxyuridine, Ki67, and doublecortin (Sahay et al. 2011a). The Nestin- CreER^T2^*Bax^fl/fl^* mouse line was kindly provided by Drs. Amar Sahay and Rene Hen and used and described previously by our group (Bermudez-Hernandez et al. 2017; Jain et al. 2019). Although Nestin-Cre-ER^T2^ mouse lines have been criticized because they can have leaky expression, the mouse line used in the present study did not (Sun et al. 2014), which we confirmed (Jain et al. 2019).

Starting at 6 weeks of age, mice were injected subcutaneously (s.c.) with tamoxifen (dose 100 mg/kg, 1/day for 5 days; Cat# T5648, Sigma-Aldrich). Tamoxifen was administered from a stock solution (20 mg/ml in corn oil, containing 10% absolute alcohol; Cat# C8267, Sigma-Aldrich). Tamoxifen is light-sensitive so it was stored at 4°C in an aluminum foil-wrapped container for the duration of treatment (5 days).

### III. Pilocarpine-induced SE

Six weeks after the last dose of tamoxifen injection, mice were injected with pilocarpine to induce SE. Methods were similar to those used previously (Jain et al. 2019). On the day of pilocarpine injection, there were 2 initial injections of pre-treatments and then one injection of pilocarpine. The first injection of pre-treatments was a solution of ethosuximide (150 mg/kg of 84 mg/ml in phosphate buffered saline, s.c.; Cat# E;7138, Sigma-Aldrich). Ethosuximide was used because the background strain, C57BL6/J, is susceptible to respiratory arrest during a severe seizure and ethosuximide decreases the susceptibility (Iyengar et al. 2015). The second injection of pre-treatments was a solution of scopolamine methyl nitrate (1 mg/kg of 0.2 mg/ml in sterile 0.9% sodium chloride solution, s.c.; Cat# 2250, Sigma-Aldrich) and terbutaline hemisulfate (1 mg/kg of 0.2 mg/ml in sterile 0.9% sodium chloride solution, s.c.; Cat# T2528, Sigma-Aldrich). Scopolamine is a muscarinic cholinergic antagonist and when injected as methyl nitrate it does not cross the blood brain barrier. Therefore, scopolamine decreased peripheral cholinergic side effects of pilocarpine without interfering with central actions of pilocarpine. Terbutaline was used to keep airways patent during severe seizures, minimizing mortality. Ethosuximide had to be administered separately because it precipitates when mixed with scopolamine and terbutaline.

Thirty min after the pre-treatments, pilocarpine hydrochloride was injected (260- 280 mg/kg of 50 mg/ml in sterile 0.9% sodium chloride solution, s.c.; Cat# P6503; Sigma-Aldrich). Different doses were used because different batches of pilocarpine had different ability to elicit SE.

The severity of SE was decreased by administering the benzodiazepine diazepam (10 mg/kg of 5 mg/ml stock solution, s.c.; NDC# 0409-3213-12, Hospira, Inc.) 2 hr after pilocarpine injection. In females, diazepam was injected earlier, 40 minutes after the onset of first seizure, because in the first group of females in which diazepam was injected 2 hr after pilocarpine, there was severe brain damage. While sedated with diazepam, animals were injected with warm (31°C) lactated Ringer’s solution (s.c.; NDC# 07-893-1389, Aspen Veterinary Resources). At the end of the day, mice were injected with ethosuximide using the same dose as before pilocarpine. For the next 3 days, chow was provided that was moistened with water. The cage was placed on a heating blanket to maintain cage temperature at 31°C.

### IV. Stereotaxic surgery

#### A. General information

Mice were anesthetized by isoflurane inhalation (3% isoflurane for induction and 1.75 - 2% isoflurane for maintenance during surgery; NDC# 07-893-1389, Patterson Veterinary) and placed in a stereotaxic apparatus (David Kopf Instruments). Prior to surgery, the analgesic Buprenex (Buprenorphine hydrochloride; NDC# 1296-0757-5; Reckitt Benckheiser) was diluted in sterile saline (0.9% sodium chloride solution) to yield a 0.03 mg/ml stock solution and 0.2 mg/kg was injected s.c. During surgery, mice were placed on a heating blanket with a rectal probe for automatic maintenance of body temperature at 31°C.

#### B. Implantation of EEG electrodes

Before electrode implantation, hair over the skull was shaved and then the scalp was cleaned with 70% ethanol. A midline incision was made to expose the skull with a sterile scalpel. To implant subdural screw electrodes (0.10” stainless steel screws, Cat# 8209, Pinnacle Technology), 6 holes were drilled over the exposed skull. The coordinates were: right occipital cortex (anterior-posterior or AP −3.5 mm from Bregma, medio-lateral or ML, 2.0 mm from the midline); left frontal cortex (Lt FC, AP −0.5 mm; ML −1.5 mm); left hippocampus (AP −2.5 mm; ML −2.0 mm) and right hippocampus (AP −2.5 mm; ML 2.0 mm). An additional screw was placed over the right olfactory bulb as ground (AP 2.3 mm; ML 1.8 mm) and another screw over the cerebellum at the midline as reference (relative to Lambda: AP −1.5 mm; ML −0.5 mm). Here, “ground” refers to the earth ground and “reference” refers to the reference for all 4 screw electrode recordings (Moyer et al. 2017). An 8-pin connector (Cat# ED85100-ND, Digi-Key Corporation) was placed over the skull and secured with dental cement (Cat# 51459, Dental Cement Kit; Stoelting Co.).

After surgery, mice were injected with 50 ml/kg warm (31°C) lactated Ringer’s solution (s.c.; NDC# 09355000476, Aspen Veterinary Resources). Mice were housed a clean cage on a heating blanket for 24 hr. Moistened food pellets were placed at the base of the cage to encourage food intake. Afterwards mice were housed individually because group housing leads to disturbance of the implant by other mice in the cage.

### V. Continuous Video-EEG recording and analysis

#### A. Video-EEG recording

Mice were allowed 3 weeks to recover from surgery. During this time, mice were housed in the room where video-EEG equipment are placed so that mice could acclimate to the recording environment. To record video-EEG, the pin connector on the head of the mouse was attached to a preamplifier (Cat# 8406, Pinnacle Technology) which was attached to a commutator (Cat# 8408, Mouse Swivel/Commutator, 4- channel, Pinnacle Technology) to allow freedom of movement. Signals were acquired at a 500 Hz sampling rate, and band-pass filtered at 1-100 Hz using Sirenia Acquisition software (https://www.pinnaclet.com, RRID:SCR_016183). Video was captured with a high-intensity infrared LED camera (Cat# PE-605EH, Pecham) and was synchronized to the EEG record.

To monitor pilocarpine-induced SE, video-EEG was recorded for hour before ad 24 hr after pilocarpine injection. Approximately 5-6 weeks after pilocarpine-induced SE, video-EEG was recorded to measure spontaneous recurrent seizures. Video-EEG was recorded continuously for 3 weeks.

#### B. Video-EEG analysis

EEG was analyzed offline with Sirenia Seizure Pro, V2.0.7 (Pinnacle Technology, RRID:SCR_016184). A seizure was defined as a period of rhythmic (>3 Hz) deflections that were >2x the standard deviation of baseline mean and lasted at least 10 sec (Jain et al. 2019). Seizures were rated as convulsive if an electrographic seizure was accompanied by a behavioral convulsion (observed by video playback), defined as stages 3-5 using the Racine scale (Racine 1972) where stage 3 is unilateral forelimb clonus, stage 4 is bilateral forelimb clonus with rearing, and stage 5 is bilateral forelimb clonus followed by rearing and falling. A seizure was defined as non-convulsive when there was electrographic evidence of a seizure but there were no stage 3-5 behavior.

SE was defined as continuous seizures for >5 min (Chen and Wasterlain 2006) and EEG amplitude in all 4 channels >3x the baseline mean. For mice without EEG, SE was defined by stage 3-5 seizures that did not stop with a resumption of normal behavior. Often stage 3-5 seizures heralded the onset of SE and then occurred intermittently for hours. In between convulsive behavior mice had twitching of their body, typically in a prone position.

SE duration was defined in light of the fact that the EEG did not return to normal after the initial period of intense activity. Instead, intermittent spiking occurred for at least 24 hrs, as we previously described (Jain et al. 2019) and has been described by others (Mazzuferi et al. 2012; Bumanglag and Sloviter 2018; Smith et al. 2018). We therefore chose a definition that captured the initial, intense activity. We defined the end of this time as the point when the amplitude of the EEG deflections were reduced to 50% or less of the peak deflections during the initial hour of SE. Specifically, we selected the time after the onset of SE when the EEG amplitude in at least 3 channels had dropped to approximately 2 times the amplitude of the EEG during the first hour of SE, and remained depressed for at least 10 min (Fig S2 in (Jain et al. 2019). Thus, the duration of SE was defined as the time between the onset and this definition of the “end” of SE.

To access the severity of chronic seizures, frequency and duration of seizures were measured during the 3 weeks of EEG recording. Inter-cluster interval was defined as the maximum number of days between two clusters.

### VI. Tissue processing

#### A. Perfusion-fixation and sectioning

Mice were perfused after video-EEG recording. To perfuse, mice were deeply anesthetized by isoflurane inhalation followed by urethane (250 mg/kg of 250 mg/ml in 0.9% sodium chloride, intraperitoneal, i.p.; Cat#U2500; Sigma-Aldrich). After loss of a reflex to a tail pinch, and loss of a righting reflex, consistent with deep anesthesia, the heart cavity was opened, and a 25-gauge needle inserted into the heart, followed by perfusion with 10 ml saline (0.9% sodium chloride in double-distilled water (ddH_2_O) using a peristaltic pump (Minipuls 1; Gilson) followed by 30 ml of cold (4°C) 4% paraformaldehyde (PFA; Cat# 19210, Electron Microscopy Sciences) in 0.1 M phosphate buffer (PB; pH 7.4). The brains were removed immediately, hemisected, and post-fixed for at least 24 hr in 4% PFA at 4°C. After post-fixation, one hemisphere was cut in the coronal plane and the other in the horizontal plane (50 μm-thick sections) using a vibratome (Cat# TPI-3000, Vibratome Co.). Sections were collected sequentially to select sections that were from similar septotemporal levels. For dorsal hippocampus, coronal sections were selected every 300 μm starting at the first section where the DG blades are fully formed (between AP −1.94 and −2.06 mm). Horizontal sections were chosen every 300 μm starting from the temporal pole at the place where the GCL is clearly defined (between DV 0.84 and 1.08 mm). This scheme is diagrammed and described in more detail in prior studies (Moretto et al. 2017).

#### B. Doublecortin

##### 1) Procedures for staining

Doublecortin (DCX), a microtubule-associated protein (Gleeson et al. 1999), was used to identify immature adult-born neurons (Brown et al. 2003; Couillard-Despres et al. 2005), and was stained after antigen retrieval (Botterill et al. 2015). First, free floating sections were washed in 0.1 M Tris buffer (TB, 3 × 5 min). Sections were then incubated in sodium citrate (Cat# S4641, Sigma-Aldrich) buffer (2.94 mg/ml in ddH_2_O, pH 6.0 adjusted with HCl) in a preheated water bath at 85°C for 30 min. Sections were washed with 0.1 M TB (3 × 5 min), blocked in 5% goat serum (Cat# S-1000, RRID:AB_2336615, Vector Laboratories) in 0.1 M TB with 0.5% (v/v) Triton X-100 (Cat# X-100, Sigma-Aldrich) and 1% (w/v) bovine serum albumin for 1 hr. Next, sections were incubated overnight with primary antibody (1:1000 diluted in blocking serum, monoclonal anti-rabbit DCX; Cat#4604S, Cell Signaling Technology) on a shaker (Model# BDRAA115S, Stovall Life Science Inc.) at room temperature.

On the next day, sections were washed in 0.1 M TB (3 × 5 min), treated with 2.5% hydrogen peroxide (Cat# 216763, Sigma-Aldrich) for 30 min to block endogenous peroxide, and washed with 0.1 M TB (3 × 5 min). Next, sections were incubated in secondary antibody (biotinylated goat anti-rabbit IgG, 1:500, Vector Laboratories) for 1 hr in 0.1 M TB, followed by washes with 0.1 M TB (3 × 5 min). Sections were then incubated in avidin-biotin complex (1:500 in 0.1 M Tris buffer; Cat# PK-6100, Vector) for 2 hr, washed in 0.1 M TB (1 × 5 min) and then in 0.175 M sodium acetate (14.36 mg/ml in ddH_2_O, pH 6.8, adjusted with glacial acetic acid, 2 × 5 min; Cat# S8750, Sigma-Aldrich). Sections were reacted in 0.5 mg/ml 3, 3′-diaminobenzidine (DAB; Cat# D5905, Sigma-Aldrich) with 40 µg/ml ammonium chloride (Cat# A4514, Sigma-Aldrich), 3 µg/ml glucose oxidase (Cat# G2133, Sigma-Aldrich), 2 mg/ml (D+)-glucose (Cat# G5767, Sigma-Aldrich) and 25 mg/ml ammonium nickel sulfate (Cat# A1827, Sigma-Aldrich) in 0.175 M sodium acetate. Sections were washed in 0.175 M sodium acetate (2 × 5 min) and 0.1 M TB (5 min), mounted on gelatin-coated slides (1% bovine gelatin; Cat# G9391, Sigma-Aldrich), and dried overnight at room temperature.

On the next day, sections were dehydrated with increasing concentrations of ethanol, cleared in Xylene (Cat# 534-56, Sigma-Aldrich), and coverslipped with Permount (Cat# 17986-01, Electron Microscopy Sciences). Sections were viewed with a brightfield microscope (Model BX51; Olympus of America) and images were captured with a digital camera (Model Infinity3-6URC, Teledyne Lumenera).

##### 2) DCX Analysis

DCX was quantified by first defining a region of interest (ROI) that included the adult-born cells and the majority of their DCX-labeled dendrites: the SGZ, GCL, and inner molecular layer. The SGZ was defined as a region that extended from the GCL into the hilus for a width of 100 µm because this region included the vast majority of the DCX immunoreactivity. The inner molecular layer was defined as the 100 μm immediately above the GCL. Next, a threshold was selected where DCX-immunoreactive (ir) cells were above, but the background was below threshold, as described in more detail elsewhere (Jain et al. 2019).

This measurement is referred to as area fraction in the Results and expressed as a percent. For a given animal, the area fraction was determined for 3 coronal sections in the dorsal hippocampus between AP -1.94 to -2.06 mm and 3-4 horizontal sections in the ventral hippocampus between DV 0.84 to 1.08 mm, with sections spaced 300 μm apart. These area fractions were averaged so that a mean area fraction was defined for each animal. For these and other analyses described below, the investigator was blinded.

#### C. Prox-1

##### 1) Procedures for staining

In normal rodent adult brain, prospero homeobox 1 (Prox1) is expressed in the GCs (Pleasure et al. 2000) and in the hilus (Bermudez-Hernandez et al. 2017). To stain for Prox1, free-floating sections were washed in 0.1 M TB pH 7.4, 3×5 min). Sections were then incubated in 0.1 M TB with 0.25%Triton X-100 for 30 min followed by a 10 min-long wash in 0.1 M TB with 0.1%Triton X-100 (referred to as Tris A). Next, sections were treated with 1% hydrogen peroxide in Tris A for 5 min followed by a 5 min-long wash in Tris A. Sections were blocked in 10% normal horse serum (Cat# S-2000, RRID:AB_2336617, Vector) in Tris A for 1 hr followed by a 10 min-long wash in Tris A and then 0.1 M TB with 0.1% Triton X-100 and 0.005% bovine serum albumin (referred to as Tris B). Next, sections were incubated overnight with primary antibody (goat anti-human Prox1 polyclonal antibody, 1:2,000 diluted in Tris B, R and D systems) rotated on a shaker (described above) at room temperature.

On the next day, sections were washed in Tris A then in Tris B (5 min each).

Sections were then incubated in secondary antibody (biotinylated anti-goat IgG made in horse, 1:500, Vector Laboratories, see Table 2) for 1 hr in Tris B, followed by a wash with Tris A (5 min) and then Tris B (5 min), blocked in avidin-biotin complex (1:500 in Tris B) for 2 hr, and washed in 0.1 M TB (3 × 5 min). Sections were reacted in 0.5 mg/ml 3, 3′-diaminobenzidine (DAB) with 40 µg/ml ammonium chloride, 3 µg/ml glucose oxidase, 2 mg/ml (D+)-glucose and 5mM nickel chloride (Cat# N6136, Sigma-Aldrich) in 0.1 M TB. Sections were washed in 0.1 M TB (3 x 5 min), mounted on 1% gelatin-coated slides and dried overnight at room temperature. On the next day, sections were dehydrated, cleared, and coverslipped (as described above). Sections were viewed and images were captured as DCX above.

##### 2) Prox1 Analysis

Prox1 was quantified in the hilus, defined based on zone 4 of Amaral (Amaral 1978). The definition of Amaral was modified to exclude 20 μm below the GCL (Winawer et al. 2007). The GCL boundary was defined as the location where GCs stopped being contiguous. Practically that meant there was no GC with more than a cell body width of cell-free space around it. A cell body width was 10 μm (Claiborne et al. 1990; Amaral et al. 2007).

CA3c was included in the ROI but hilar Prox1 cells have not been detected in the CA3c layer (Scharfman et al. 2000; Winawer et al. 2007). However, there are rare GCs in CA3 according to one study (Szabadics et al. 2010).

In ImageJ, a ROI was traced in the image taken at 20x magnification and then a threshold was selected where Prox1-immunoreactivity was above the background threshold (Jain et al. 2019). Then Prox1 cells were counted using the Analyzed particle plugin where a particle with an area ≥ 10 μm^2^ was counted. The following criteria were used to define a hilar Prox1-ir cell (Bermudez-Hernandez et al. 2017): (1) the hilar cell had sufficient Prox1-ir to reach a threshold equal to the average level of Prox1-ir of GCs in the adjacent GC layer, (2) All hilar Prox-ir cells were complete, i.e., not cut at the edge of the ROI. When hilar Prox1-ir cells were in clusters, although not many (typically 2–3 per 50 μm section), cells were counted manually. For each animal 3 coronal sections in the dorsal hippocampus and 3-4 horizontal sections in the ventral hippocampus, with sections spaced 300 μm apart were chosen.

#### D. Immunofluorescence

##### 1) Procedures for staining

Free floating sections were washed (3×5 min) in 0.1 M TB followed by a 10-min long wash in Tris A and another 10 min-long wash in Tris B. Sections were incubated in blocking solution (5% normal goat serum or donkey serum in Tris B) for 1 hr at room temperature. Next, primary antibodies for anti-rabbit GluR2/3, anti-goat Prox1, anti-rabbit SOM and anti-mouse parvalbumin (Table 1) were diluted in blocking solution and sections were incubated for 48 hr at 4°C. For SOM labelling, antigen retrieval was used. Prior to the blocking step, sections were incubated in sodium citrate buffer (2.94 mg/ml in ddH_2_O, pH 6.0 adjusted with HCl) in a preheated water bath at 85°C for 30 min.

**Table 1.**
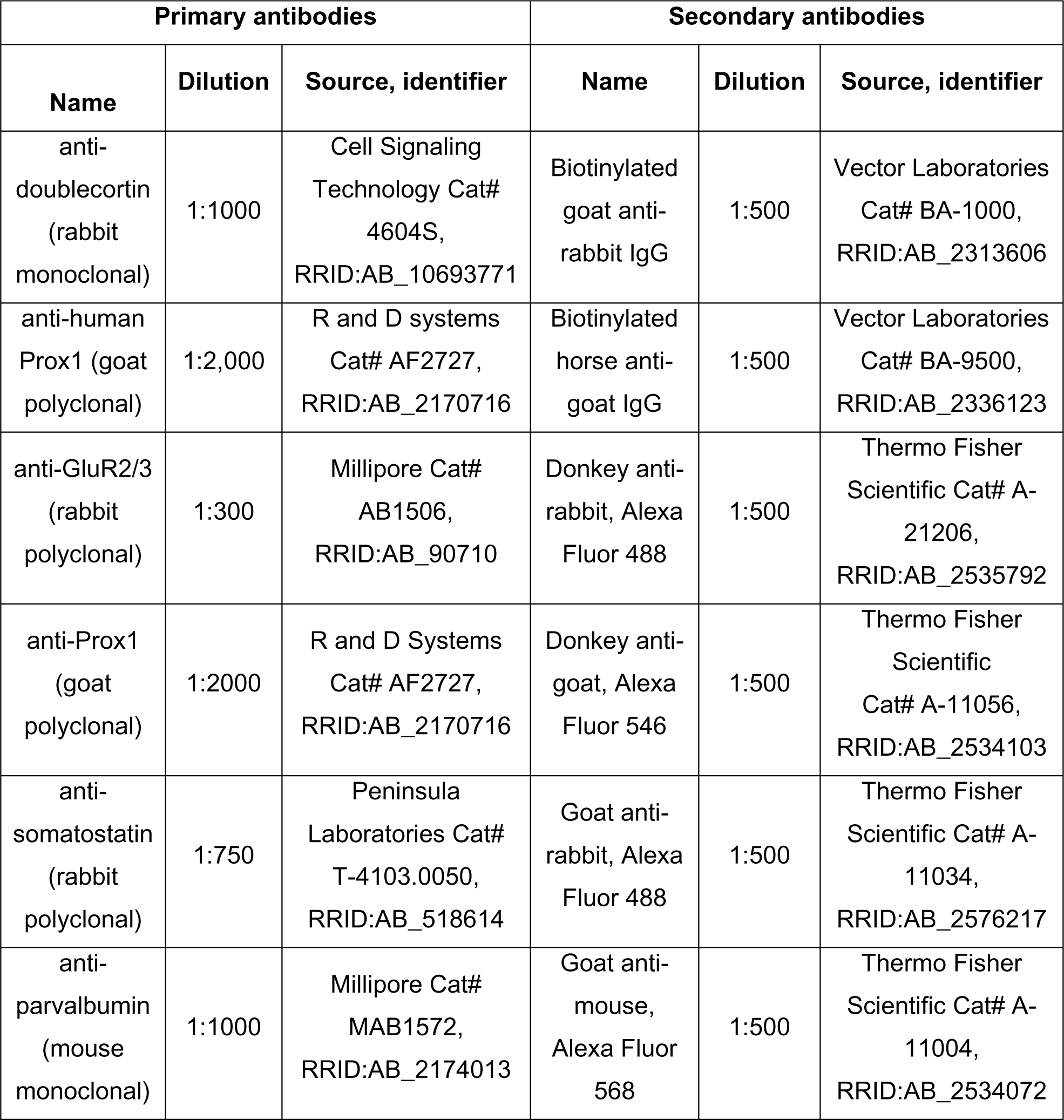
Primary antibodies.

Next, sections were washed in Tris A and Tris B (10 min each) followed by 2 hr-long incubation in secondary antibody (1:500 in Tris B, see Table 2). Sections were washed in 0.1 M TB (3 x 5 min), and coverslipped with Citifluor^TM^ AF1 mounting solution (Cat# 17970-25, Vector Labs). Images were captured on a confocal microscope (Model LSM 510 Meta; Carl Zeiss Microimaging).

##### 2) Procedures for analysis

GluR2/3-, SOM- and parvalbumin- ir cells in the hilus and SGZ were counted from 3 dorsal and 3 ventral sections. Sections were viewed at 40x of the confocal microscope for manual counts. Because ectopic GCs express GluR2/3, sections were co-labelled with Prox1. All co-labelled cells were considered as ectopic and excluded from the GluR2/3- ir cell counting to measure mossy cells.

#### E. Fluorojade-C

##### 1) Procedures for staining

Fluorojade-C (FJ) is a fluorescent dye that is the “gold standard” to stain degenerating neurons (Schmued and Hopkins 2000; Schmued et al. 2005). First, sections were mounted on gelatin-coated slides (1% porcine gelatin in ddH2O; Cat# G1890, Sigma-Aldrich) and dried on a hot plate at 50–55°C for 1 hr. Then slides were placed in a staining rack and immersed in a 100% ethanol solution for 5 min, then in 70% ethanol for 2 min, followed by a 1 min wash in ddH2O.

Slides were then incubated in 0.06% potassium permanganate (Cat# P-279, Fisher Scientific) solution for 10 min on a shaker (described above) with gentle agitation, followed by washes in ddH2O (2 × 1 min). Slides were then incubated for 20 min in a 0.0002% solution of FJ in ddH2O with 0.1% acetic acid in the dark. The stock solution of FJ was 0.01% in ddH2O and was stored at 4°C for up to 3 months. To prepare a working solution, 6 ml of stock solution was added to 294 mL of 0.1% acetic acid (Cat# UN2789, Fisher Scientific) in ddH2O and used within 10 min of preparation. Slides were subsequently protected from direct light. They were washed in ddH2O (3 × 1 min) and dried overnight at room temperature. On the next day, slides were cleared in Xylene (2 × 3 min) and coverslipped with DPX mounting medium (Cat# 44581, Sigma-Aldrich). Sections were photographed with an epifluorescence microscope (Model BX51; Olympus of America) and images were captured with a digital camera (Model Infinity3-6URC, Teledyne Lumenera).

##### 2) Procedures for analysis

We measured the FJ in the cell layers of CA1 and CA3. Manual counting of FJ- positive (FJ+) cells was not possible in these cell layers because there could be so many FJ+ cells that were overlapping. Instead, FJ staining in cell layers was quantified by first outlining the cell layer as a ROI at 10× magnification in ImageJ as before (Jain et al. 2019).

To outline the CA1 cell layer, the border with CA2 was defined as the point where the cell layer changed width, a sudden change that could be appreciated by the background in FJ-stained sections and confirmed by cresyl violet-stained sections. The border of CA1 and the subiculum was defined as the location where the normally compact CA1 cell layer suddenly dispersed. To outline CA3, the border with CA2 and CA3 was defined by the point where stratum lucidum of CA3 terminated. This location was distinct in its background in FJ-stained sections. The border of CA3 and the hilus was defined according to zone 4 of Amaral (Amaral 1978). This location was also possible to detect in FJ-stained sections because the background in the hilus was relatively dark compared to area CA3.

After defining ROIs, a threshold fluorescence level was selected so that all cells that had very bright immunofluorescence were above threshold but other cells that were similar in fluorescence to background staining were not (Iyengar et al. 2015; Jain et al. 2019). ImageJ was then used to calculate the area within the ROI and this measurement is referred to as area fraction in the Results and expressed as a percent. For a given animal, the area fraction was determined for three coronal sections in the dorsal hippocampus between AP −1.94 and −2.06 mm and 3–4 horizontal sections in the ventral hippocampus between DV 0.84 and 1.08 mm, with sections spaced 300 μm apart. These area fractions were averaged so that a mean area fraction was defined for each animal.

### VI. Statistical Analysis

Data are presented as the mean ± standard error of the mean (SEM). Statistical analyses were performed using GraphPad Prism Software (https://www.graphpad.com/ <u>scientific-software/prism/</u>, RRID: SCR_002798). Statistical significance was set at p < 0.05. Robust regression and Outlier removal (ROUT) method was used to remove outliers with ROUT coefficient Q set at 1%. Parametric tests were used when data fit a normal distribution, determined by the D’Agostino and Pearson or Shapiro-Wilk’s normality tests, and there was homoscedasticity of variance (confirmed by a F-test). A Student’s unpaired two-tailed t-test was used to assess differences between two groups. A Welch’s test was used instead of a Student’s t-test when there was heteroscedasity of variance. One-way Analysis of Variance (ANOVA), two-way ANOVA, and three-way ANOVA were performed when there were multiple groups and were followed by Bonferroni’s multiple comparison post-hoc test (Bonferroni’s test). The main factors for two-way ANOVA were genotype and sex; region was added as another factor for three-way ANOVA. Interaction between factors is reported in the Results if it was significant. A Fisher’s exact test was used for comparing proportions of binary data (yes/no). Pearson’s Correlation was used to assess the association between 2 variables.

For data that did not follow a normal distribution, typically some data had a 0 value. In these cases, non-parametric tests were selected. The Mann-Whitney *U* test was used to compare two groups, and a Kruskal-Wallis test followed by post-hoc Dunn’s test was used for multiple groups comparison.

## Acknowledgements

This study was supported by NIH R01 NS081203, NIH R37 NS126529 and the New York State Office of Mental Health.

## SUPPLEMENTARY FIGURES

**Fig. 1.- Supplemental Fig. 1.**
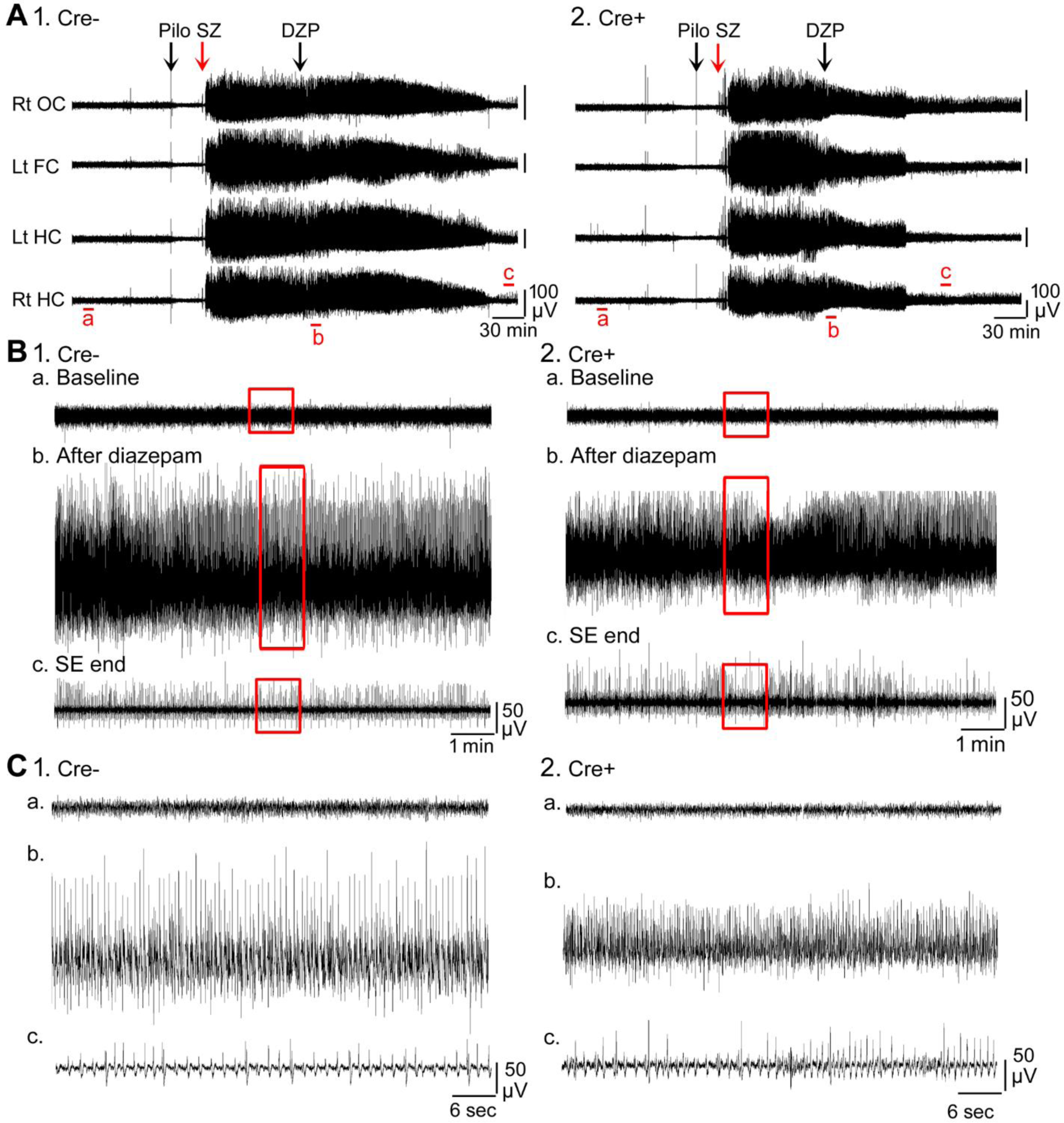
Examples of EEG during SE. A. Representative examples of a 10 hr-long EEG recording are shown for Cre- (1) and Cre+ (2) mice. These records are the same as in Fig. 1. **B.** 10 min-long EEG recording segments from the left hippocampus A are shown with higher temporal gain. 1. Cre- mouse. a. Part of the baseline is shown. The area surrounded by the red box is expanded in C1a. b. The time when DZP was injected is shown. The area surrounded by the red box is expanded in C1b. c. The time following the seizure is shown. The area surrounded by the red box is expanded in C1c. 2. Cre+ mouse. a. Part of the baseline is shown. The area surrounded by the red box is expanded in C2a. Note the baselines are similar in the two mice, suggesting no effect of genotype. b. The time when DZP was injected is shown. The area surrounded by the red box is expanded in C2b. Note there was a reduction in EEG amplitude in the Cre+ mouse. c. The time at the end of SE is shown. The area surrounded by the red box is expanded in C2c. Note that these two mice were similar after SE ended. **C.** The areas surrounded by the red boxes in B are expanded. The traces are 1 min- long.

**Fig. 1.- Supplemental Fig. 2.**
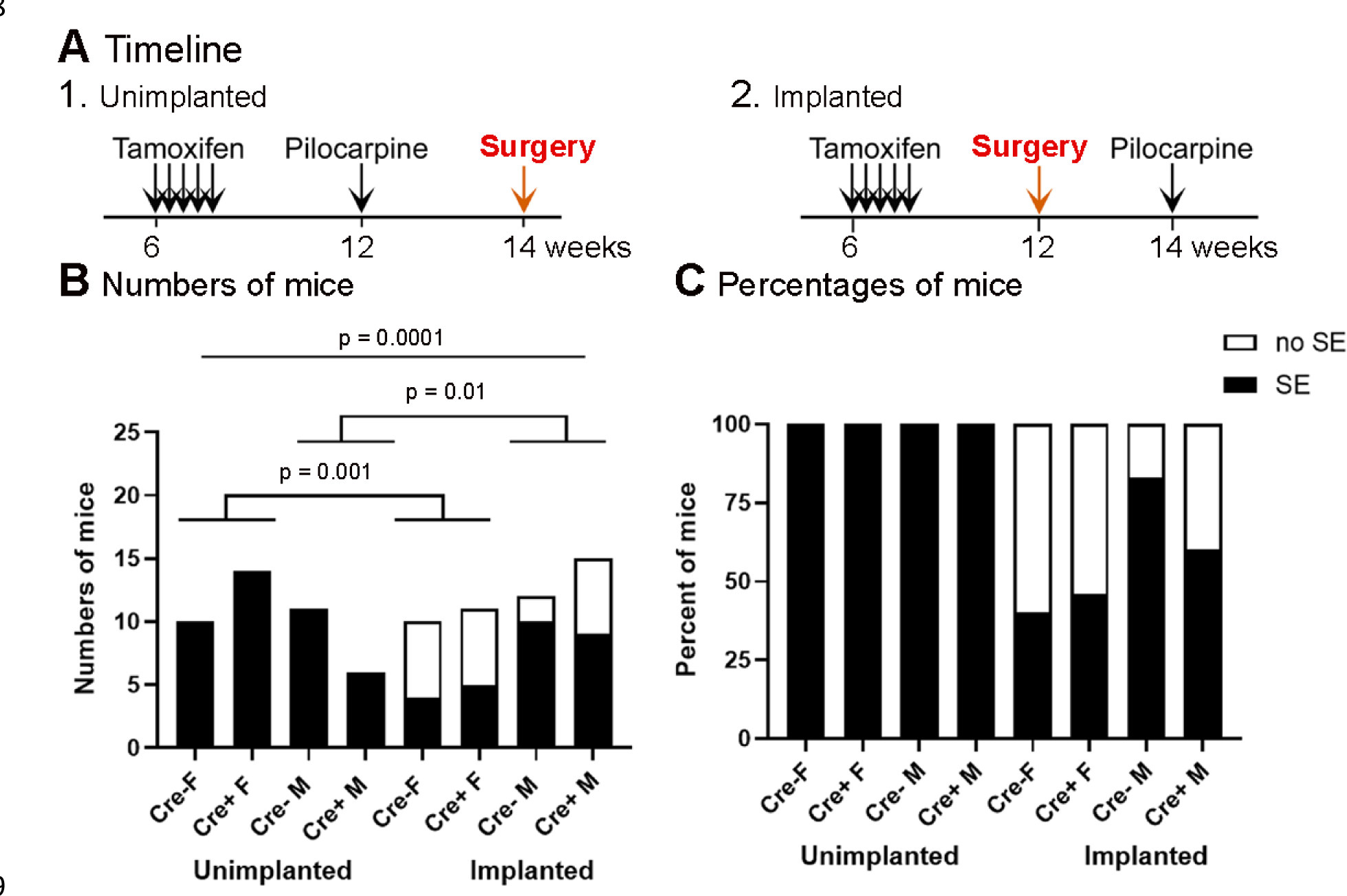
Incidence of SE in the unimplanted and implanted mice. A. The experimental timelines of pilocarpine injection and surgery. 1. Unimplanted mice. 2. Implanted mice. Only the timing of surgery with respect to pilocarpine injection was different. **B.** The incidence of SE in unimplanted and implanted mice is shown based on numbers of mice. The incidence of SE was significantly higher in the unimplanted mice (Fisher’s exact test, p<0.0001). Genotype had no effect on the incidence of SE. **C.** The incidence of SE is shown as percentages.

**Fig. 1.- Supplemental Fig. 3.**
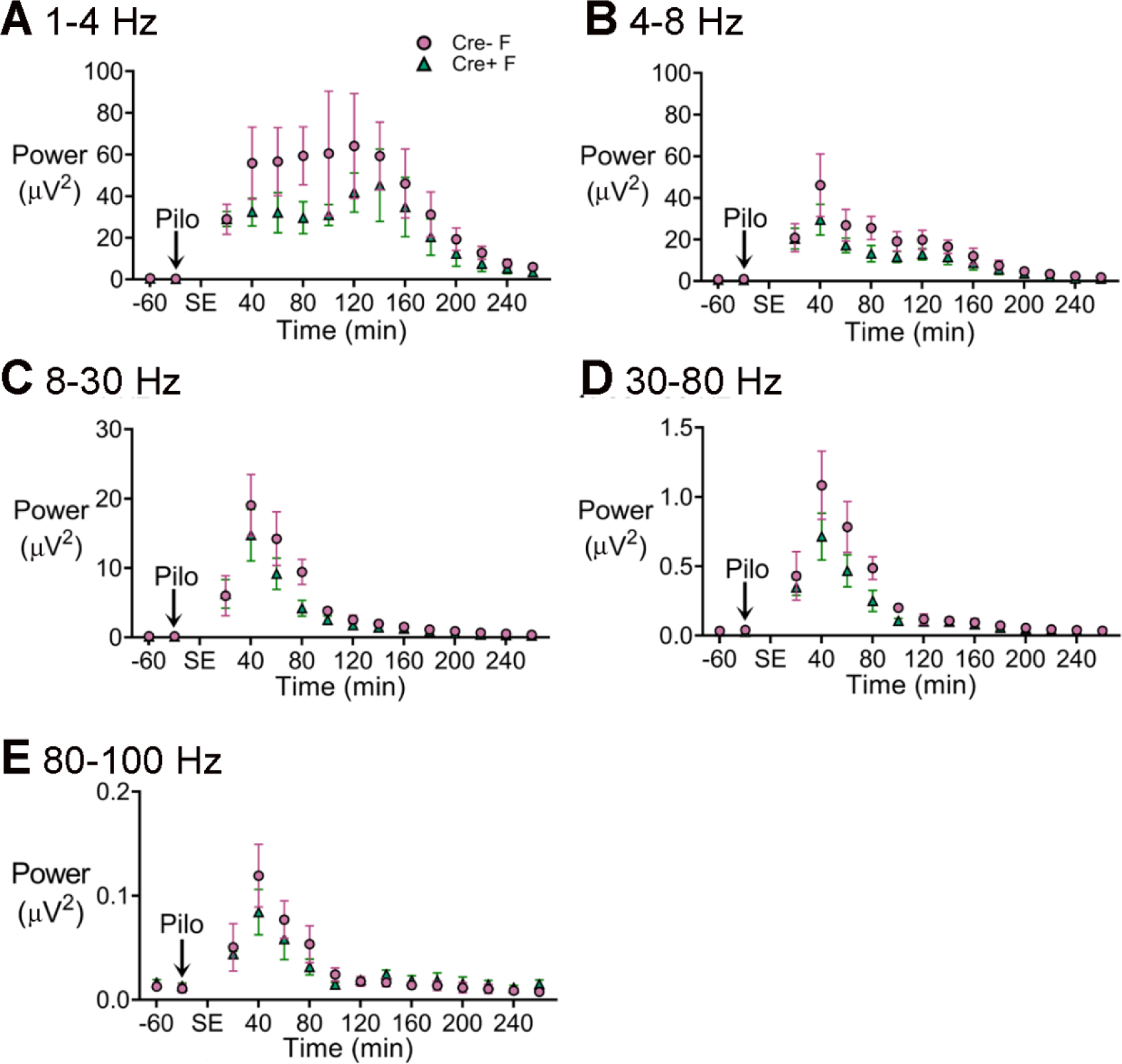
Power during SE in Cre+ female and Cre- female mice. A. Power was calculated for consecutive 20 min-long bins before and during SE. Power in the 1-4 Hz band was decreased during SE in female Cre+ mice (blue triangles) relative to Cre- mice (red circles) but it was not statistically significant. **B.** Power in the 4-8 Hz band. **C.** Power in 8-30 Hz range. **D.** Power in 30-80 Hz range **E.** Power between 80 and 100 Hz.

**Fig. 1- Supplemental Fig. 4.**
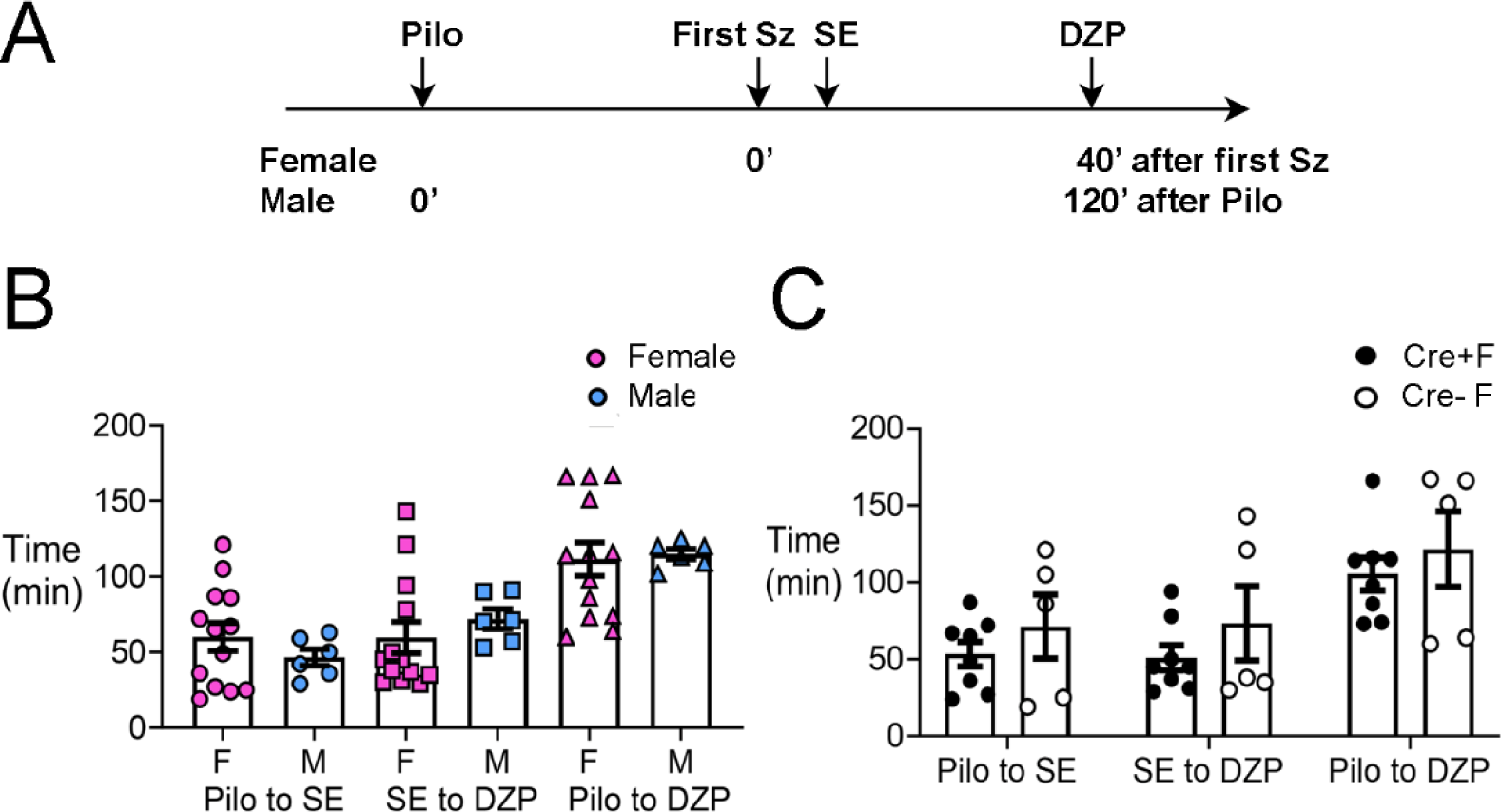
The latency to SE, interval between SE and diazepam administration, and interval between pilocarpine and diazepam injections were not significantly different in experimental groups. A. A timeline of experimental procedures is shown for the day of pilocarpine-induced SE. Pilocarpine was injected and the first seizure stage 3 or greater was noted. The onset of SE was noted also. For females, diazepam (DZP) was injected 45 min after the first seizure (Sz). Males were administered DZP 2 hrs after pilocarpine (Pilo). The reason for the difference is that it made the latencies to SE, interval between SE and DZP injection, and interval between pilocarpine and DZP injections similar. **B.** The mean ± SEM is shown for the time from pilocarpine to SE, SE to DZP injection, and pilocarpine to DZP injection. There were no sex differences: a two-way ANOVA with sex and type of measurement as main factors showed no effect of sex (F(1,45)=0004, p=0.949). However, there were differences between the types of measurements (F(2,45)=11.89, p<0.0001). **C.** When Cre+ and Cre- females were compared, there was no effect of genotype (F(1,33)=2.33, p=0.136) but there was a significant effect of the type of measurement (F(2,33)=7.66, p=0.002).

**Fig. 2.- Supplemental Fig. 1.**
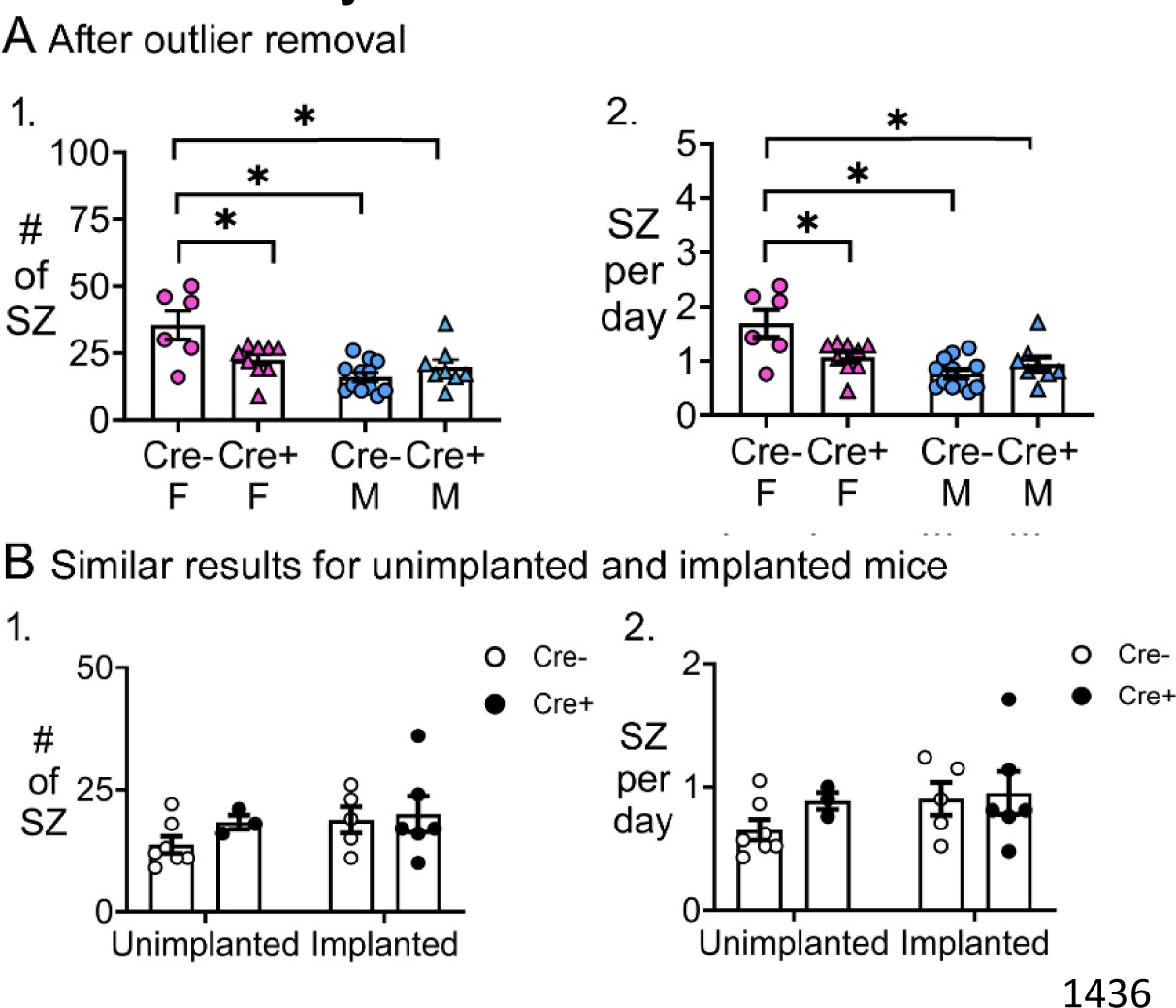
Additional analyses of chronic seizures. Similar results after outlier removal. 1. A two-way ANOVA with sex and genotype as factors showed a significant effect of sex on the # of seizures (F(1,31)=16.04, p=0.0004) and an interaction (F1,31)=9.20, p=0.005) with no main effect of genotype (F(1,31)=1.75, p=0.107). Cre- females had significantly more seizures than all other groups (post-hoc tests, Cre- females vs. Cre+ females, p=0.020; vs. Cre- males, p=0.0002; vs. Cre+ males, p=0.005). Cre- males and Cre+ males were not different (p=0.722). 2. A two-way ANOVA with sex and genotype as factors showed a significant effect of sex (F(1,31)=15.84, p=0.0004) on seizure frequency and an interaction (F(1,31)=9.16, p=0.005) although no main effect of genotype (F(1,31)=2.72, p=0.109). Cre- females had significantly more seizures than all other groups (post-hoc tests, Cre- females vs. Cre+ females, p=0.021; vs. Cre- males, p=0.0002; vs. Cre+ males, 0.005). Cre- males were not different from Cre+ males (p=0.720). **B.** Results were independent of the time when EEG electrodes were implanted. 1. The number of chronic seizures were similar between mice that were implanted before and after SE by two-way ANOVA with implant status and genotype as factors (implant status, F(1,17)=1.33, p=0.265; genotype, F(1,17)=0.88, p=0.334). Sexes were pooled. 2. The frequency of seizures was similar between mice implanted before or after SE (implant status, F(1,17)=1.27, p=0.276; genotype, F(1,17)=1.00, p=0.330). Sexes were pooled.

**Fig. 2.- Supplemental Fig. 2.**
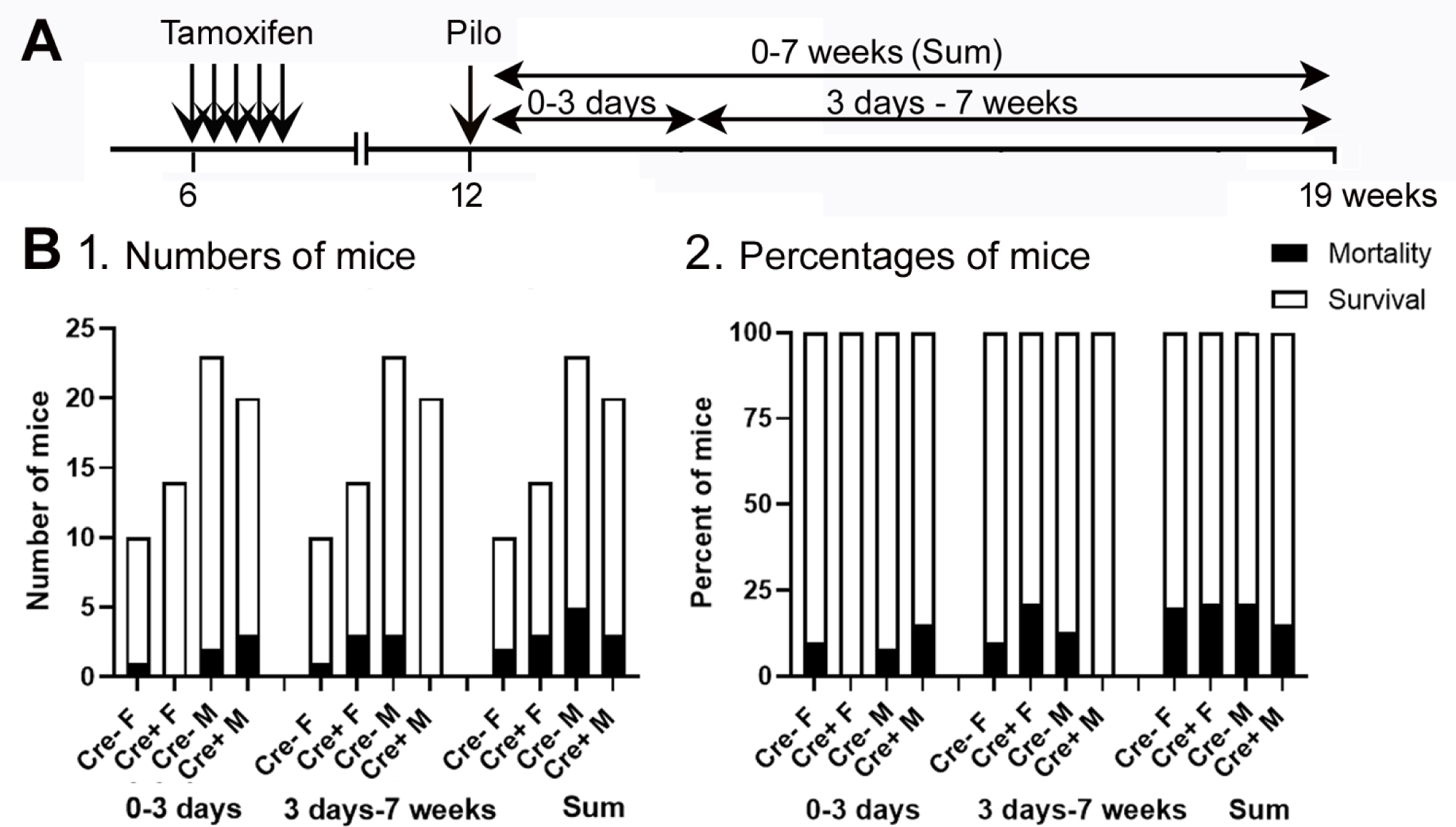
**Mortality was not significantly affected by genotype or sex.** A. The timeline of measurements of mortality is shown. Mice were categorized as dying during SE or within 3 days of SE (0-3 days), 3 days to 7 weeks, or both (Sum). Mice are included whether they were implanted with EEG electrodes before SE or implanted 3 weeks after SE. 1. The numbers of mice that died were not significantly different between genotypes or sexes (Fisher’s Exact test, all p>0.05). 2. The percent of mice that died is shown.

**Fig. 5.- Supplemental Fig. 1.**
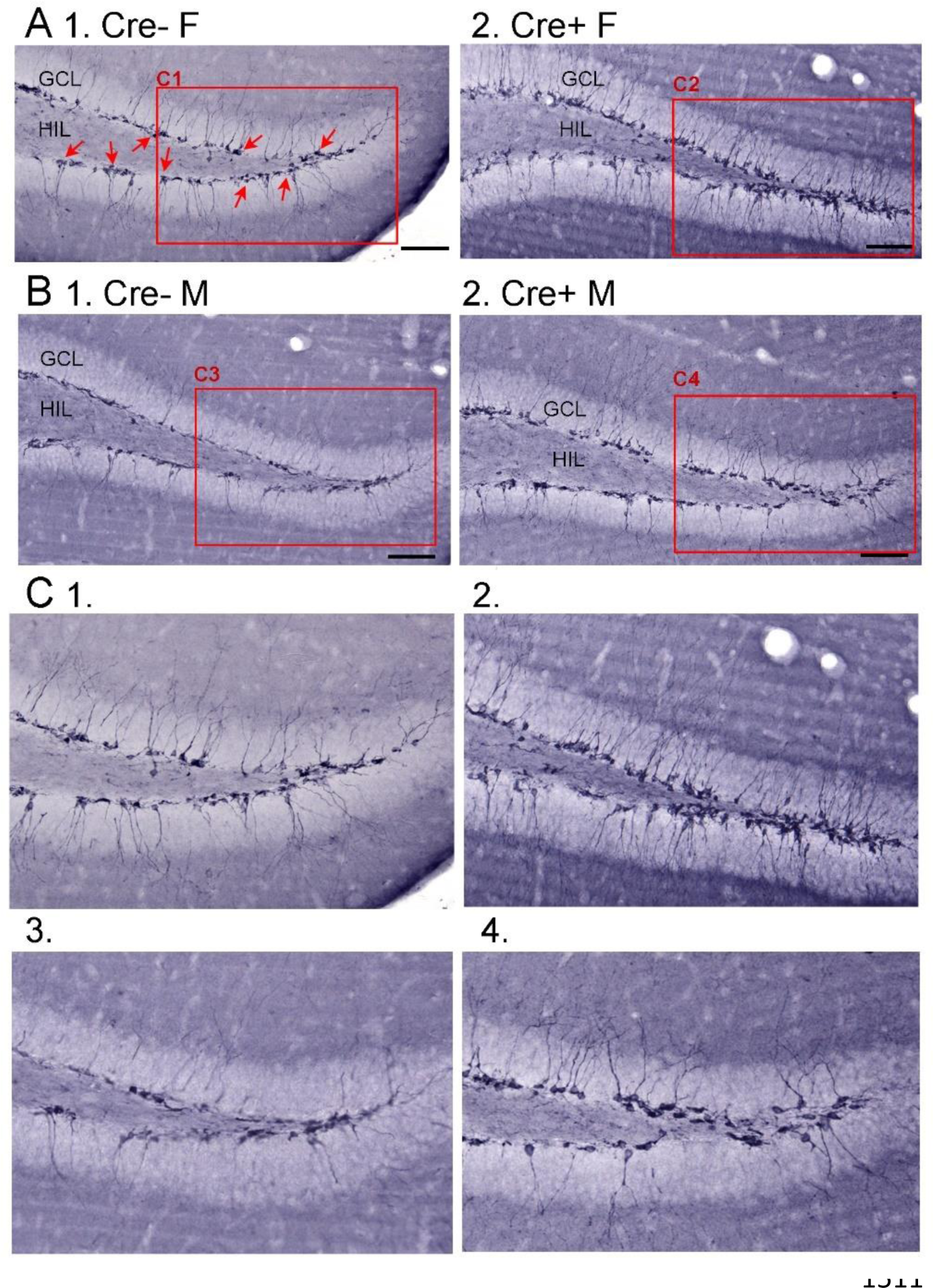
DCX in Cre- and Cre+ mice before SE. A. Mice were administered tamoxifen for 5 days at 6 weeks of age as for other experiments. Six weeks after tamoxifen, mice were perfusion-fixed and staining was conducted using an antibody to DCX. 1. Representative image from a Cre- female (F) of dorsal dentate gyrus in coronal section shows many DCX-ir cells in the SGZ and GCL (red arrows). The area surrounded by the red box is expanded in C1. Calibration, 100 µm. 2. Example of DCX-ir from a Cre+ female mouse shows more DCX-ir, reflecting more immature neurons. The area surrounded by the red box is expanded in C2. Calibration, 100 µm. 3. Example from a Cre- male (M). The area surrounded by the red box is expanded in C3. 4. Example of a Cre+ M showing more DCX-ir than the Cre- M. The area surrounded by the red box is expanded in C4. Calibration, 100 µm. **B.** Expanded insets from A-B. 1. Cre- F; 2. Cre+ F, 3. Cre- M, 4. Cre+ M.

**Fig. 7.- Supplemental Fig. 1.**
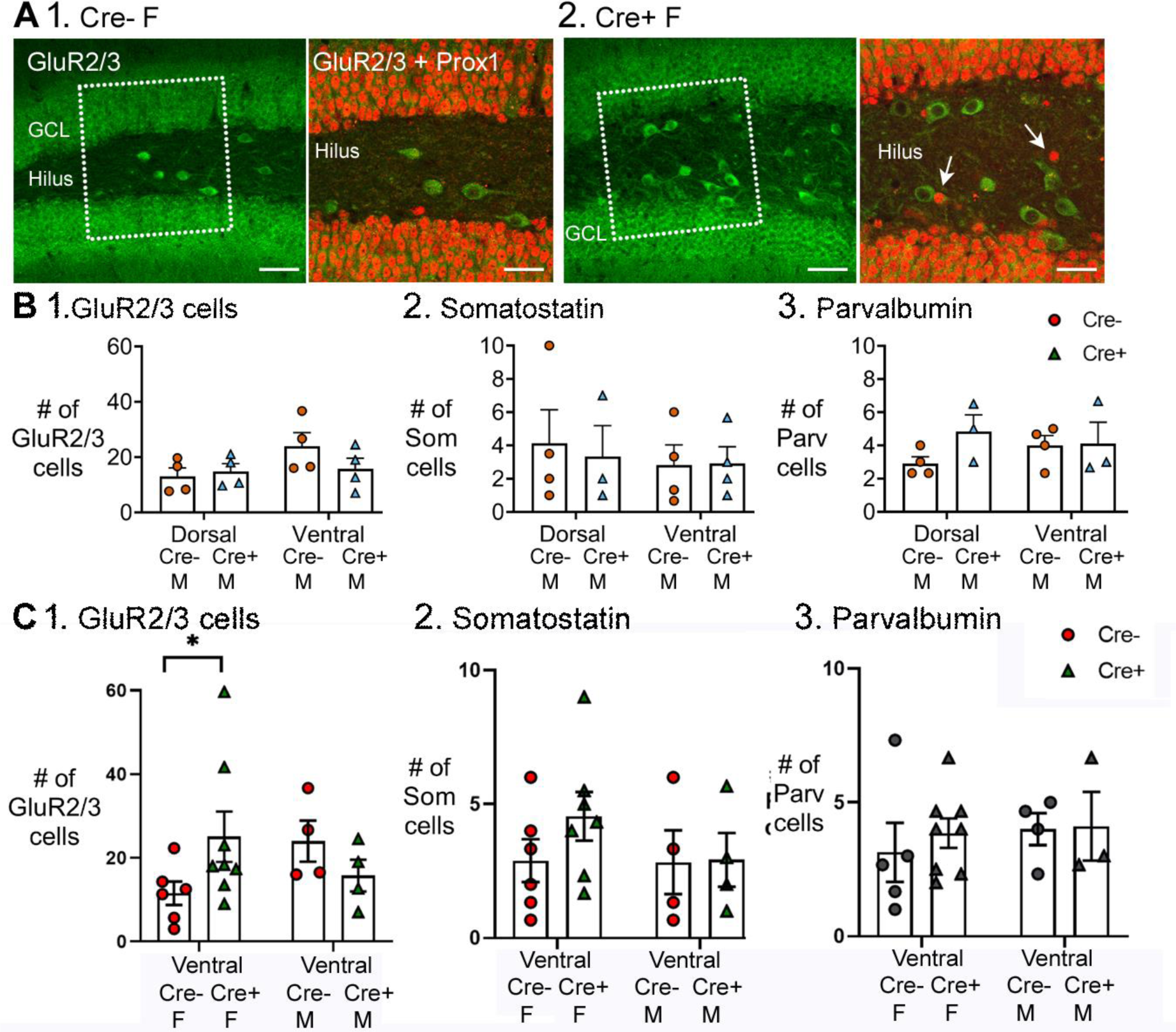
Additional analyses of GluR2/3, SOM and parvalbumin-expressing cells. A. GluR2/3 hilar cells lacked Prox1 expression. 1. Cre- female mouse. Left: several GluR2/3+ cells (green) are located in the hilus within the box (marked by dotted white lines). Calibration, 70 µm. Right: The area within the box in the left panel is expanded. The merged image of GluR2/3+ (green) and Prox1+ (red) cells shows no double labeling. Calibration, 40 µm. 2. Cre + female mouse. Similar results are shown as for the Cre- mouse. White arrows mark ectopic GCs. Calibrations are the same as for the Cre- mouse. **B.** A comparison of dorsal and ventral measurements for Cre- and Cre+ male mice show no significant genotype effects. 1. GluR2/3. A two-way ANOVA showed no effect of dorsal or ventral location (F(1,13)=3.38; p=0.089) or genotype (F(1,13)=1.158; p=0.302). 2. SOM. A two-way ANOVA showed no effect of dorsal or ventral location (F(1,10)=0.172; p=0.687) or genotype (F(1,10)=0.014; p=0.908). 3. Parvalbumin. A two-way ANOVA showed no effect of dorsal or ventral location (F(1,13)=0.358; p=0.560) or genotype (F(1,13)=1.068 p=0.320). **C.** A comparison of ventral measurements for both Cre- and Cre+ female and male mice. 1. There were significantly more GluR2/3+ hilar cells in Cre+ female mice compared to Cre- female mice, like the dorsal hippocampus (Fig. 7). Thus, GluR2/3+ hilar cells were spared in Cre+ females in dorsal and ventral hippocampus. A two-way ANOVA showed o effect of sex (F(1,18)=0.744; p=0.400) or genotype F(1,18)=0.386; p=0.542) but there was a significant interaction F(1,18)=5.433; p=0.0316, and post-hoc tests showed that Cre+ females had significantly more GluR2/3+ cells than Cre- females (p=0.045). 2. There were no significant differences among groups for SOM+ cells. Thus, there was an effect in dorsal (Fig. 7) but not ventral hippocampus. Thus, SOM cells were spared in Cre+ females dorsally but not ventrally. A two-way ANOVA showed no effect of sex (F(1,17)=0.718; p=0.408) or genotype F (1, 17)=0.769; P=0.393). There were no significant differences in numbers of parvalbumin+ cells, like dorsal hippocampus (Fig. 7). Thus, parvalbumin cells were similar regardless of genotype in dorsal and ventral hippocampus. A two-way ANOVA showed no significant effect of sex (F(1,16)=0.401; p=0.536) or genotype (F(1,16)=0.221; p=0.645).

## REFERENCES

1. Acsady L, Kamondi A, Sik A, Freund T and Buzsaki G (1998) Gabaergic cells are the major postsynaptic targets of mossy fibers in the rat hippocampus. J Neurosci. 18, 3386–3403.

2. Adlaf EW, Vaden RJ, Niver AJ, Manuel AF, Onyilo VC, Araujo MT, Dieni CV, Vo HT, King GD, Wadiche JI and Overstreet-Wadiche L (2017) Adult-born neurons modify excitatory synaptic transmission to existing neurons. Elife. 6, e19886.

3. Althaus AL, Moore SJ, Zhang H, Du X, Murphy GG and Parent JM (2019) Altered synaptic drive onto birthdated dentate granule cells in experimental temporal lobe epilepsy. J Neurosci. 39, 7604–7614.

4. Althaus AL, Zhang H and Parent JM (2016) Axonal plasticity of age-defined dentate granule cells in a rat model of mesial temporal lobe epilepsy. Neurobiol Dis. 86, 187–196.

5. Altman J 2011. The discovery of adult mammalian neurogenesis. In: Seki T, Sawamoto K, Parent JM & Alvarez-Buylla A (eds.) Neurogenesis in the adult brain 1: Neurobiology. Springer.

6. Altman J and Das GD (1965) Autoradiographic and histological evidence of postnatal hippocampal neurogenesis in rats. J Comp Neurol. 124, 319–335.

7. Amaral DG (1978) A Golgi study of cell types in the hilar region of the hippocampus in the rat. J Comp Neurol. 182, 851–914.

8. Amaral DG, Scharfman HE and Lavenex P (2007) The dentate gyrus: Fundamental neuroanatomical organization (dentate gyrus for dummies). Prog Brain Res. 163, 3–22.

9. Andre V, Marescaux C, Nehlig A and Fritschy JM (2001) Alterations of hippocampal gabaergic system contribute to development of spontaneous recurrent seizures in the rat lithium-pilocarpine model of temporal lobe epilepsy. Hippocampus. 11, 452–468.

10. Ash AM, Regele-Blasco E, Seib DR, Chahley E, Skelton PD, Luikart BW and Snyder JS (2023) Adult-born neurons inhibit developmentally-born neurons during spatial learning. Neurobiol Learn Mem. 198, 107710.

11. Ben-Ari Y and Holmes GL (2006) Effects of seizures on developmental processes in the immature brain. Lancet Neurol. 5, 1055–1063.

12. Bermudez-Hernandez K, Lu YL, Moretto J, Jain S, LaFrancois JJ, Duffy AM and Scharfman HE (2017) Hilar granule cells of the mouse dentate gyrus: Effects of age, septotemporal location, strain, and selective deletion of the proapoptotic gene *Bax*. Brain Struct Funct. 222, 3147–3161.

13. Bolay H, Berman NE and Akcali D (2011) Sex-related differences in animal models of migraine headache. Headache. 51, 891–904.

14. Boldrini M, Fulmore CA, Tartt AN, Simeon LR, Pavlova I, Poposka V, Rosoklija GB, Stankov A, Arango V, Dwork AJ, Hen R and Mann JJ (2018) Human hippocampal neurogenesis persists throughout aging. Cell Stem Cell. 22, 589–599 e585.

15. Botterill JJ, Brymer KJ, Caruncho HJ and Kalynchuk LE (2015) Aberrant hippocampal neurogenesis after limbic kindling: Relationship to BDNF and hippocampal-dependent memory. Epilepsy Behav. 47, 83–92.

16. Botterill JJ, Lu YL, LaFrancois JJ, Bernstein HL, Alcantara-Gonzalez D, Jain S, Leary P and Scharfman HE (2019) An excitatory and epileptogenic effect of dentate gyrus mossy cells in a mouse model of epilepsy. Cell Rep. 29, 2875–2889 e2876.

17. Brown JP, Couillard-Despres S, Cooper-Kuhn CM, Winkler J, Aigner L and Kuhn HG (2003) Transient expression of doublecortin during adult neurogenesis. J Comp Neurol. 467, 1–10.

18. Bui AD, Nguyen TM, Limouse C, Kim HK, Szabo GG, Felong S, Maroso M and Soltesz I (2018) Dentate gyrus mossy cells control spontaneous convulsive seizures and spatial memory. Science. 359, 787–790.

19. Bumanglag AV and Sloviter RS (2018) No latency to dentate granule cell epileptogenesis in experimental temporal lobe epilepsy with hippocampal sclerosis. Epilepsia. 59, 2019–2034.

20. Cameron HA, Woolley CS, McEwen BS and Gould E (1993) Differentiation of newly born neurons and glia in the dentate gyrus of the adult rat. Neuroscience. 56, 337–344.

21. Cavalheiro EA, Santos NF and Priel MR (1996) The pilocarpine model of epilepsy in mice. Epilepsia. 37, 1015–1019.

22. Cavazos JE, Das I and Sutula TP (1994) Neuronal loss induced in limbic pathways by kindling: Evidence for induction of hippocampal sclerosis by repeated brief seizures. J Neurosci. 14, 3106–3121.

23. Cavazos JE and Sutula TP (1990) Progressive neuronal loss induced by kindling: A possible mechanism for mossy fiber synaptic reorganization and hippocampal sclerosis. Brain Res. 527, 1–6.

24. Chancey JH, Poulsen DJ, Wadiche JI and Overstreet-Wadiche L (2014) Hilar mossy cells provide the first glutamatergic synapses to adult-born dentate granule cells. J Neurosci. 34, 2349–2354.

25. Chen JW and Wasterlain CG (2006) Status epilepticus: Pathophysiology and management in adults. Lancet Neurol. 5, 246–256.

26. Cho KO, Lybrand ZR, Ito N, Brulet R, Tafacory F, Zhang L, Good L, Ure K, Kernie SG, Birnbaum SG, Scharfman HE, Eisch AJ and Hsieh J (2015) Aberrant hippocampal neurogenesis contributes to epilepsy and associated cognitive decline. Nat Commun. 6.

27. Choi SH, Bylykbashi E, Chatila ZK, Lee SW, Pulli B, Clemenson GD, Kim E, Rompala A, Oram MK, Asselin C, Aronson J, Zhang C, Miller SJ, Lesinski A, Chen JW, Kim DY, van Praag H, Spiegelman BM, Gage FH and Tanzi RE (2018) Combined adult neurogenesis and BDNF mimic exercise effects on cognition in an Alzheimer’s mouse model. Science. 361.

28. Choi SH and Tanzi RE (2019) Is Alzheimer’s disease a neurogenesis disorder? Cell Stem Cell. 25, 7–8.

29. Claiborne BJ, Amaral DG and Cowan WM (1990) Quantitative, three-dimensional analysis of granule cell dendrites in the rat dentate gyrus. J Comp Neurol. 302, 206–219.

30. Clelland CD, Choi M, Romberg C, Clemenson GD, Jr., Fragniere A, Tyers P, Jessberger S, Saksida LM, Barker RA, Gage FH and Bussey TJ (2009) A functional role for adult hippocampal neurogenesis in spatial pattern separation. Science. 325, 210–213.

31. Couillard-Despres S, Winner B, Schaubeck S, Aigner R, Vroemen M, Weidner N, Bogdahn U, Winkler J, Kuhn HG and Aigner L (2005) Doublecortin expression levels in adult brain reflect neurogenesis. Eur J Neurosci. 21, 1–14.

32. Coulter DA and Carlson GC (2007) Functional regulation of the dentate gyrus by GABA-mediated inhibition. Prog Brain Res. 163, 235–243.

33. de Lanerolle NC, Kim JH, Robbins RJ and Spencer DD (1989) Hippocampal interneuron loss and plasticity in human temporal lobe epilepsy. Brain Res. 495, 387–395.

34. Dingledine R, Varvel NH and Dudek FE (2014) When and how do seizures kill neurons, and is cell death relevant to epileptogenesis? Adv Exp Med Biol. 813, 109–122.

35. Drew LJ, Kheirbek MA, Luna VM, Denny CA, Cloidt MA, Wu MV, Jain S, Scharfman HE and Hen R (2016) Activation of local inhibitory circuits in the dentate gyrus by adult-born neurons. Hippocampus. 26, 763–778.

36. Dudek FE and Staley KJ 2012. The time course and circuit mechanisms of acquired epileptogenesis. In: Noebels J, Avoli M, Rogawski M, Olsen R & Delgado-Escueta A (eds.) Jasper’s basic mechanisms of the epilepsies. 4th ed. Bethesda, MD: Oxford University Press.

37. Eikermann-Haerter K, Dilekoz E, Kudo C, Savitz SI, Waeber C, Baum MJ, Ferrari MD, van den Maagdenberg AM, Moskowitz MA and Ayata C (2009) Genetic and hormonal factors modulate spreading depression and transient hemiparesis in mouse models of familial hemiplegic migraine type 1. J Clin Invest. 119, 99–109.

38. Falconer MA, Serafetinides EA and Corsellis JA (1964) Etiology and pathogenesis of temporal lobe epilepsy. Arch Neurol. 10, 233–248.

39. Forger NG, Rosen GJ, Waters EM, Jacob D, Simerly RB and de Vries GJ (2004) Deletion of Bax eliminates sex differences in the mouse forebrain. Proc Natl Acad Sci U S A. 101, 13666–13671.

40. Freund TF, Ylinen A, Miettinen R, Pitkänen A, Lahtinen H, Baimbridge KG and Riekkinen PJ (1992) Pattern of neuronal death in the rat hippocampus after status epilepticus. Relationship to calcium binding protein content and ischemic vulnerability. Brain Res Bull. 28, 27–38.

41. Gage F, Kempermann G and Song H 2008. Adult neurogenesis, New York, Cold Spring Harbor Laboratory Press.

42. Galeeva A, Treuter E, Tomarev S and Pelto-Huikko M (2007) A prospero-related homeobox gene prox-1 is expressed during postnatal brain development as well as in the adult rodent brain. Neuroscience. 146, 604–616.

43. Galichet C, Guillemot F and Parras CM (2008) Neurogenin 2 has an essential role in development of the dentate gyrus. Development. 135, 2031–2041.

44. Gleeson JG, Lin PT, Flanagan LA and Walsh CA (1999) Doublecortin is a microtubule-associated protein and is expressed widely by migrating neurons. Neuron. 23, 257–271.

45. Goffin K, Nissinen J, Van Laere K and Pitkanen A (2007) Cyclicity of spontaneous recurrent seizures in pilocarpine model of temporal lobe epilepsy in rat. Exp Neurol. 205, 501–505.

46. Han ZS, Buhl EH, Lorinczi Z and Somogyi P (1993) A high degree of spatial selectivity in the axonal and dendritic domains of physiologically identified local-circuit neurons in the dentate gyrus of the rat hippocampus. Eur J Neurosci. 5, 395–410.

47. Hartings JA, Shuttleworth CW, Kirov SA, Ayata C, Hinzman JM, Foreman B, Andrew RD, Boutelle MG, Brennan KC, Carlson AP, Dahlem MA, Drenckhahn C, Dohmen C, Fabricius M, Farkas E, Feuerstein D, Graf R, Helbok R, Lauritzen M, Major S, Oliveira-Ferreira AI, Richter F, Rosenthal ES, Sakowitz OW, Sanchez- Porras R, Santos E, Scholl M, Strong AJ, Urbach A, Westover MB, Winkler MK, Witte OW, Woitzik J and Dreier JP (2017) The continuum of spreading depolarizations in acute cortical lesion development: Examining leao’s legacy. J Cereb Blood Flow Metab. 37, 1571–1594.

48. Haut SR (2015) Seizure clusters: Characteristics and treatment. Curr Opin Neurol. 28, 143–150.

49. Henshall DC and Meldrum BS 2012. Cell death and survival mechanisms after single and repeated brief seizures. In: Noebels JL, Avoli M, Rogawski MA, Olsen RW & Delgado-Escueta AV (eds.) Jasper’s basic mechanisms of the epilepsies. 4th ed. Bethesda, MD: Oxford University Press.

50. Henze DA, Urban NN and Barrionuevo G (2000) The multifarious hippocampal mossy fiber pathway: A review. Neuroscience. 98, 407–427.

51. Herman ST (2002) Epilepsy after brain insult: Targeting epileptogenesis. Neurology. 59, S21–26.

52. Herreras O and Makarova J (2020) Mechanisms of the negative potential associated with leao’s spreading depolarization: A history of brain electrogenesis. J Cereb Blood Flow Metab. 40, 1934–1952.

53. Hester MS and Danzer SC (2013) Accumulation of abnormal adult-generated hippocampal granule cells predicts seizure frequency and severity. J Neurosci. 33, 8926–8936.

54. Hosford BE, Liska JP and Danzer SC (2016) Ablation of newly generated hippocampal granule cells has disease-modifying effects in epilepsy. J Neurosci. 36, 11013–11023.

55. Hsu D (2007) The dentate gyrus as a filter or gate: A look back and a look ahead. Prog Brain Res. 163, 601–613.

56. Huusko N, Romer C, Ndode-Ekane XE, Lukasiuk K and Pitkanen A (2015) Loss of hippocampal interneurons and epileptogenesis: A comparison of two animal models of acquired epilepsy. Brain Struct Funct. 220, 153–191.

57. Ikrar T, Guo N, He K, Besnard A, Levinson S, Hill A, Lee HK, Hen R, Xu X and Sahay A (2013) Adult neurogenesis modifies excitability of the dentate gyrus. Front Neural Circuits. 7, 204–235.

58. Iwano T, Masuda A, Kiyonari H, Enomoto H and Matsuzaki F (2012) Prox1 postmitotically defines dentate gyrus cells by specifying granule cell identity over CA3 pyramidal cell fate in the hippocampus. Development. 139, 3051–3062.

59. Iyengar SS, LaFrancois JJ, Friedman D, Drew LJ, Denny CA, Burghardt NS, Wu MV, Hsieh J, Hen R and Scharfman HE (2015) Suppression of adult neurogenesis increases the acute effects of kainic acid. Exp Neurol. 264, 135–149.

60. Jafarpour S, Hirsch LJ, Gaínza-Lein M, Kellinghaus C and Detyniecki K (2019) Seizure cluster: Definition, prevalence, consequences, and management. Seizure. 68, 9–15.

61. Jain S, LaFrancois JJ, Botterill JJ, Alcantara-Gonzalez D and Scharfman HE (2019) Adult neurogenesis in the mouse dentate gyrus protects the hippocampus from neuronal injury following severe seizures. Hippocampus. 29, 683–709.

62. Jakubs K, Nanobashvili A, Bonde S, Ekdahl CT, Kokaia Z, Kokaia M and Lindvall O (2006) Environment matters: Synaptic properties of neurons born in the epileptic adult brain develop to reduce excitability. Neuron. 52, 1047–1059.

63. Jung KH, Chu K, Lee ST, Kim J, Sinn DI, Kim JM, Park DK, Lee JJ, Kim SU, Kim M, Lee SK and Roh JK (2006) Cyclooxygenase-2 inhibitor, celecoxib, inhibits the altered hippocampal neurogenesis with attenuation of spontaneous recurrent seizures following pilocarpine-induced status epilepticus. Neurobiol Dis. 23, 237–246.

64. Kaplan MS and Hinds JW (1977) Neurogenesis in the adult rat: Electron microscopic analysis of light radioautographs. Science. 197, 1092–1094.

65. Kazanis I 2013. Neurogenesis in the adult mammalian brain: How much do we need, how much do we have? In: Belzung C & Wigmore P (eds.) Neurogenesis and neural plasticity. New York: Springer.

66. Kempermann G 2012. Adult hippocampal neurogenesis. In: Kempermann G (Ed.) Adult neurogenesis 2. 2 ed. New York: Oxford University Press.

67. Kempermann G, Gage FH, Aigner L, Song H, Curtis MA, Thuret S, Kuhn HG, Jessberger S, Frankland PW, Cameron HA, Gould E, Hen R, Abrous DN, Toni N, Schinder AF, Zhao X, Lucassen PJ and Frisen J (2018) Human adult neurogenesis: Evidence and remaining questions. Cell Stem Cell. 23, 25–30.

68. Kempermann G, Song H and Gage FH (2015) Neurogenesis in the adult hippocampus. Cold Spring Harb Perspect Biol. 7.

69. Kron MM, Zhang H and Parent JM (2010) The developmental stage of dentate granule cells dictates their contribution to seizure-induced plasticity. J Neurosci. 30, 2051–2059.

70. Krook-Magnuson E, Armstrong C, Bui A, Lew S, Oijala M and Soltesz I (2015) In vivo evaluation of the dentate gate theory in epilepsy. J Physiol. 593, 2379–2388.

71. Kudo C, Harriott AM, Moskowitz MA, Waeber C and Ayata C (2023) Estrogen modulation of cortical spreading depression. J Headache Pain. 24, 62.

72. Leranth C, Malcolm AJ and Frotscher M (1990) Afferent and efferent synaptic connections of somatostatin-immunoreactive neurons in the rat fascia dentata. J Comp Neurol. 295, 111–122.

73. Leranth C, Szeidemann Z, Hsu M and Buzsaki G (1996) AMPA receptors in the rat and primate hippocampus: A possible absence of glur2/3 subunits in most interneurons. Neuroscience. 70, 631–652.

74. Levesque M, Biagini G, de Curtis M, Gnatkovsky V, Pitsch J, Wang S and Avoli M (2021) The pilocarpine model of mesial temporal lobe epilepsy: Over one decade later, with more rodent species and new investigative approaches. Neurosci Biobehav Rev. 130, 274–291.

75. Lu YL and Scharfman HE (2021) New insights and methods for recording and imaging spontaneous spreading depolarizations and seizure-like events in mouse hippocampal slices. Front Cell Neurosci. 15, 761423.

76. Mathern GW, Wilson CL and Beck H 2008. Hippocampal sclerosis. In: Engel J, Pedley TA & Aicardi J (eds.) Epilepsy: A comprehensive textbook. 2nd ed. Philadelphia, PA: Lippincott Williams & Wilkins.

77. Mazzuferi M, Kumar G, Rospo C and Kaminski RM (2012) Rapid epileptogenesis in the mouse pilocarpine model: Video-EEG, pharmacokinetic and histopathological characterization. Exp Neurol. 238, 156–167.

78. McCloskey DP, Hintz TM, Pierce JP and Scharfman HE (2006) Stereological methods reveal the robust size and stability of ectopic hilar granule cells after pilocarpine-induced status epilepticus in the adult rat. Eur J Neurosci. 24, 2203–2210.

79. Moreno-Jimenez EP, Flor-Garcia M, Terreros-Roncal J, Rabano A, Cafini F, Pallas-Bazarra N, Avila J and Llorens-Martin M (2019) Adult hippocampal neurogenesis is abundant in neurologically healthy subjects and drops sharply in patients with Alzheimer’s disease. Nat Med. 25, 554–560.

80. Moretto JN, Duffy AM and Scharfman HE (2017) Acute restraint stress decreases c-fos immunoreactivity in hilar mossy cells of the adult dentate gyrus. Brain Struct Funct. 222, 2405–2419.

81. Moyer JT, Gnatkovsky V, Ono T, Otahal J, Wagenaar J, Stacey WC, Noebels J, Ikeda A, Staley K, de Curtis M, Litt B and Galanopoulou AS (2017) Standards for data acquisition and software-based analysis of in vivo electroencephalography recordings from animals. A task1-wg5 report of the aes/ilae translational task force of the ilae. Epilepsia. 58 Suppl 4, 53-67.

82. Myers CE, Bermudez-Hernandez K and Scharfman HE (2013) The influence of ectopic migration of granule cells into the hilus on dentate gyrus-CA3 function. PLoS One. 8, e68208.

83. Nakashiba T, Cushman JD, Pelkey KA, Renaudineau S, Buhl DL, McHugh TJ, Rodriguez Barrera V, Chittajallu R, Iwamoto KS, McBain CJ, Fanselow MS and Tonegawa S (2012) Young dentate granule cells mediate pattern separation, whereas old granule cells facilitate pattern completion. Cell. 149, 188–201.

84. Niibori Y, Yu TS, Epp JR, Akers KG, Josselyn SA and Frankland PW (2012) Suppression of adult neurogenesis impairs population coding of similar contexts in hippocampal CA3 region. Nat Commun. 3.

85. Paredes MF, Sorrells SF, Cebrian-Silla A, Sandoval K, Qi D, Kelley KW, James D, Mayer S, Chang J, Auguste KI, Chang EF, Gutierrez Martin AJ, Kriegstein AR, Mathern GW, Oldham MC, Huang EJ, Garcia-Verdugo JM, Yang Z and Alvarez-Buylla A (2018) Does adult neurogenesis persist in the human hippocampus? Cell Stem Cell. 23, 780-781.

86. Parent JM and Kron MM 2012. Neurogenesis and epilepsy. *In:* Noebels JL, Avoli M, Rogawski MA, Olsen RW & Delgado-Escueta AV (eds.) Jasper’s basic mechanisms of the epilepsies. Bethesda (MD): National Center for Biotechnology Information (US)

87. Copyright © 2012, Michael A Rogawski, Antonio V Delgado-Escueta, Jeffrey L Noebels, Massimo Avoli and Richard W Olsen. Parent JM and Lowenstein DH (2002) Seizure-induced neurogenesis: Are more new neurons good for an adult brain? Prog Brain Res. 135, 121–131.

88. Parent JM and Murphy GG (2008) Mechanisms and functional significance of aberrant seizure-induced hippocampal neurogenesis. Epilepsia. 49 Suppl 5, 19–25.

89. Parent JM, Yu TW, Leibowitz RT, Geschwind DH, Sloviter RS and Lowenstein DH (1997) Dentate granule cell neurogenesis is increased by seizures and contributes to aberrant network reorganization in the adult rat hippocampus. J Neurosci. 17, 3727–3738.

90. Piatti VC and Schinder AF (2018) Hippocampal mossy cells provide a fate switch for adult neural stem cells. Neuron. 99, 425–427.

91. Pierce JP, McCloskey DP and Scharfman HE (2011) Morphometry of hilar ectopic granule cells in the rat. J Comp Neurol. 519, 1196–1218.

92. Pierce JP, Punsoni M, McCloskey DP and Scharfman HE (2007) Mossy cell axon synaptic contacts on ectopic granule cells that are born following pilocarpine- induced seizures. Neurosci Lett. 422, 136–140.

93. Pleasure SJ, Collins AE and Lowenstein DH (2000) Unique expression patterns of cell fate molecules delineate sequential stages of dentate gyrus development. J Neurosci. 20, 6095–6105.

94. Racine RJ (1972) Modification of seizure activity by electrical stimulation. Ii. Motor seizure. Electroencephalogr Clin Neurophysiol. 32, 281–294.

95. Ramirez-Amaya V, Marrone DF, Gage FH, Worley PF and Barnes CA (2006) Integration of new neurons into functional neural networks. J Neurosci. 26, 12237–12241.

96. Sahay A, Scobie KN, Hill AS, O’Carroll CM, Kheirbek MA, Burghardt NS, Fenton AA, Dranovsky A and Hen R (2011a) Increasing adult hippocampal neurogenesis is sufficient to improve pattern separation. Nature. 472, 466–470.

97. Sahay A, Wilson DA and Hen R (2011b) Pattern separation: A common function for new neurons in hippocampus and olfactory bulb. Neuron. 70, 582–588.

98. Savanthrapadian S, Meyer T, Elgueta C, Booker SA, Vida I and Bartos M (2014) Synaptic properties of SOM- and CCK-expressing cells in dentate gyrus interneuron networks. J Neurosci. 34, 8197–8209.

99. Scharfman H, Goodman J and McCloskey D (2007) Ectopic granule cells of the rat dentate gyrus. Dev Neurosci. 29, 14–27.

100. Scharfman HE (1999) The role of nonprincipal cells in dentate gyrus excitability and its relevance to animal models of epilepsy and temporal lobe epilepsy. Adv Neurol. 79, 805–820.

101. Scharfman HE (2004) Functional implications of seizure-induced neurogenesis. Adv Exp Med Biol. 548, 192–212.

102. Scharfman HE, Goodman JH and Sollas AL (2000) Granule-like neurons at the hilar/CA3 border after status epilepticus and their synchrony with area CA3 pyramidal cells: Functional implications of seizure-induced neurogenesis. J Neurosci. 20, 6144–6158.

103. Scharfman HE and Hen R (2007) Neuroscience. Is more neurogenesis always better? Science. 315, 336-338.

104. Scharfman HE and MacLusky NJ (2014) Differential regulation of BDNF, synaptic plasticity and sprouting in the hippocampal mossy fiber pathway of male and female rats. Neuropharmacology. 76 Pt C, 696-708.

105. Scharfman HE and McCloskey DP (2009) Postnatal neurogenesis as a therapeutic target in temporal lobe epilepsy. Epilepsy Res. 85, 150–161.

106. Scharfman HE and Pierce JP (2012) New insights into the role of hilar ectopic granule cells in the dentate gyrus based on quantitative anatomic analysis and three- dimensional reconstruction. Epilepsia. 53 Suppl 1, 109–115.

107. Scharfman HE and Schwartzkroin PA (1990a) Consequences of prolonged afferent stimulation of the rat fascia dentata: Epileptiform activity in area CA3 of hippocampus. Neuroscience. 35, 505–517.

108. Scharfman HE and Schwartzkroin PA (1990b) Responses of cells of the rat fascia dentata to prolonged stimulation of the perforant path: Sensitivity of hilar cells and changes in granule cell excitability. Neuroscience. 35, 491–504.

109. Schmued LC and Hopkins KJ (2000) Fluoro-jade b: A high affinity fluorescent marker for the localization of neuronal degeneration. Brain Res. 874, 123–130.

110. Schmued LC, Stowers CC, Scallet AC and Xu L (2005) Fluoro-jade c results in ultra high resolution and contrast labeling of degenerating neurons. Brain Res. 1035, 24–31.

111. Scorza FA, Arida RM, Naffah-Mazzacoratti Mda G, Scerni DA, Calderazzo L and Cavalheiro EA (2009) The pilocarpine model of epilepsy: What have we learned? An Acad Bras Cienc. 81, 345–365.

112. Siegel C and McCullough LD (2011) Nad+ depletion or par polymer formation: Which plays the role of executioner in ischaemic cell death? Acta Physiol (Oxf). 203, 225–234.

113. Sisk C, Lonstein J and Gore A 2016. Critical periods during development: Hormonal influences on neurobehavioral transitions across the life span. In: Pfaff D & Volkow N (eds.) Neuroscience in the 21st Century. New York: Springer.

114. Sloviter RS (1987) Decreased hippocampal inhibition and a selective loss of interneurons in experimental epilepsy. Science. 235, 73–76.

115. Sloviter RS (1994) The functional organization of the hippocampal dentate gyrus and its relevance to the pathogenesis of temporal lobe epilepsy. Ann Neurol. 35, 640–654.

116. Sloviter RS, Zappone CA, Harvey BD, Bumanglag AV, Bender RA and Frotscher M (2003) “Dormant basket cell” hypothesis revisited: Relative vulnerabilities of dentate gyrus mossy cells and inhibitory interneurons after hippocampal status epilepticus in the rat. J Comp Neurol. 459, 44–76.

117. Smith ZZ, Benison AM, Bercum FM, Dudek FE and Barth DS (2018) Progression of convulsive and nonconvulsive seizures during epileptogenesis after pilocarpine- induced status epilepticus. J Neurophysiol. 119, 1818–1835.

118. Somjen GG (2001) Mechanisms of spreading depression and hypoxic spreading depression-like depolarization. Physiol Rev. 81, 1065–1096.

119. Sorrells SF, Paredes MF, Cebrian-Silla A, Sandoval K, Qi D, Kelley KW, James D, Mayer S, Chang J, Auguste KI, Chang EF, Gutierrez AJ, Kriegstein AR, Mathern GW, Oldham MC, Huang EJ, Garcia-Verdugo JM, Yang Z and Alvarez-Buylla A (2018) Human hippocampal neurogenesis drops sharply in children to undetectable levels in adults. Nature. 555, 377–381.

120. Ssentongo P, Robuccio AE, Thuku G, Sim DG, Nabi A, Bahari F, Shanmugasundaram B, Billard MW, Geronimo A, Short KW, Drew PJ, Baccon J, Weinstein SL, Gilliam FG, Stoute JA, Chinchilli VM, Read AF, Gluckman BJ and Schiff SJ (2017) A murine model to study epilepsy and sudep induced by malaria infection. Sci Rep. 7, 43652.

121. Steiner B, Zurborg S, Horster H, Fabel K and Kempermann G (2008) Differential 24 h responsiveness of prox1-expressing precursor cells in adult hippocampal neurogenesis to physical activity, environmental enrichment, and kainic acid- induced seizures. Neuroscience. 154, 521–529.

122. Sun C, Mtchedlishvili Z, Bertram EH, Erisir A and Kapur J (2007) Selective loss of dentate hilar interneurons contributes to reduced synaptic inhibition of granule cells in an electrical stimulation-based animal model of temporal lobe epilepsy. J Comp Neurol. 500, 876–893.

123. Sun MY, Yetman MJ, Lee TC, Chen Y and Jankowsky JL (2014) Specificity and efficiency of reporter expression in adult neural progenitors vary substantially among nestin-creer(t2) lines. J Comp Neurol. 522, 1191–1208.

124. Sun W, Winseck A, Vinsant S, Park OH, Kim H and Oppenheim RW (2004) Programmed cell death of adult-generated hippocampal neurons is mediated by the proapoptotic gene *Bax*. J Neurosci. 24, 11205–11213.

125. Szabadics J, Varga C, Brunner J, Chen K and Soltesz I (2010) Granule cells in the CA3 area. J Neurosci. 30, 8296–8307.

126. Tartt AN, Fulmore CA, Liu Y, Rosoklija GB, Dwork AJ, Arango V, Hen R, Mann JJ and Boldrini M (2018) Considerations for assessing the extent of hippocampal neurogenesis in the adult and aging human brain. Cell Stem Cell. 23, 782–783.

127. Taupin P 2006. Adult neurogenesis and neural stem cells in mammals, New York, Nova Science Publishers, Inc.

128. Tobin MK, Musaraca K, Disouky A, Shetti A, Bheri A, Honer WG, Kim N, Dawe RJ, Bennett DA, Arfanakis K and Lazarov O (2019) Human hippocampal neurogenesis persists in aged adults and Alzheimer’s disease patients. Cell Stem Cell. 24, 974–982 e973.

129. Tronel S, Belnoue L, Grosjean N, Revest JM, Piazza PV, Koehl M and Abrous DN (2012) Adult-born neurons are necessary for extended contextual discrimination. Hippocampus. 22, 292–298.

130. van Vliet EA, Aronica E, Tolner EA, Lopes da Silva FH and Gorter JA (2004) Progression of temporal lobe epilepsy in the rat is associated with immunocytochemical changes in inhibitory interneurons in specific regions of the hippocampal formation. Exp Neurol. 187, 367–379.

131. Whitebirch AC, LaFrancois JJ, Jain S, Leary P, Santoro B, Siegelbaum SA and Scharfman HE (2022) Enhanced excitability of the hippocampal CA2 region and its contribution to seizure activity in a mouse model of temporal lobe epilepsy. Neuron. 110, 3121-3138 e3128.

132. Winawer MR, Makarenko N, McCloskey DP, Hintz TM, Nair N, Palmer AA and Scharfman HE (2007) Acute and chronic responses to the convulsant pilocarpine in dba/2j and a/j mice. Neuroscience. 149, 465–475.

133. Zhan RZ, Timofeeva O and Nadler JV (2010) High ratio of synaptic excitation to synaptic inhibition in hilar ectopic granule cells of pilocarpine-treated rats. J Neurophysiol. 104, 3293–3304.

134. Zhou QG, Nemes AD, Lee D, Ro EJ, Zhang J, Nowacki AS, Dymecki SM, Najm IM and Suh H (2019) Chemogenetic silencing of hippocampal neurons suppresses epileptic neural circuits. J Clin Invest. 129, 310–323.

